# Pluripotency factors enhance the firing efficiency of late DNA replication origins in mouse embryonic stem cells

**DOI:** 10.1101/2025.01.07.631761

**Authors:** Eddie Rodriguez-Carballo, Vasilis S. Dionellis, Sotirios G. Ntallis, Lilia Bernasconi, Ezgi Keskin, Thanos D. Halazonetis

**Affiliations:** Department of Molecular and Cellular Biology, University of Geneva, Switzerland; IFOM ETS, the AIRC Institute of Molecular Oncology, Milan, Italy

## Abstract

DNA replication initiates at genomic loci known as origins, which fire according to a spatiotemporal program, still not yet well-defined in mammals. Here, we investigated whether pluripotent mouse embryonic stem cells (mESCs), which proliferate rapidly and have very short G1 phases, have a different spatiotemporal origin firing program than differentiated cells. Using EdU-seq combined with various cell cycle synchronization methods, we identified DNA replication initiation zones (IZs) in mESCs, mesenchymal stem cells and mouse embryo fibroblasts (MEFs). Similar profiles of IZs were present in the early-replicating genomic domains in all cell types. Uniquely in mESCs, we observed, within 1-2 hours of entry into S phase, origin firing at IZs that mapped to the mid and late-replicating genomic domains. This change in the replication timing program, which did not lead to shortening of the length of S phase, was driven by pluripotency transcription factors, notably OCT4, as documented by examining mESCs in which OCT4 expression was transiently suppressed. We propose that pluripotency factors in mESCs bind to IZs at mid and late-replicating genomic domains to enhance chromatin accessibility, which in turn leads to changes in the replication timing program. These results provide an example of how transcription factors control DNA replication in a cell type-specific manner.

## INTRODUCTION

In mammals, genome duplication starts in a coordinated fashion at thousands of loci, the origins of replication. Origin firing takes place in a controlled-stochastic manner, meaning that a given cell activates only certain origins among a defined available set (Hu and Stillman, 2023). Molecularly, any given origin will first be licensed, following a sequence of events that starts with binding of ORC onto chromatin and recruitment of a double-hexamer MCM helicase complex. In a next step, firing of licensed origins is stimulated by DDK and CDK protein kinases, which facilitate the assembly of firing factors and replisome subunits, including TRESLIN, MTBP, TOPBP1, MCM10, CDC45 and GINS (Marchal et al., 2019a). The replication forks proceed bidirectionally, until they meet incoming forks, thus ensuring complete replication of the genome.

The genome can be split in replication timing (RT) domains, depending on when these domains are replicated during S phase. The RT program is established in G1 and shows variations between different cell types, although it shares some general features (Hiratani et al., 2008; Rausch et al., 2020; Rhind and Gilbert, 2013; Rivera-Mulia et al., 2015). Early RT domains correspond to gene-rich regions, often actively transcribed, decorated with open chromatin marks and belonging to the so-called topological A compartment in Hi-C maps, whereas late RT domains correspond to poorly-transcribed, heterochromatic regions, belonging to the B compartment in Hi-C maps (Gilbert et al., 2010; Pope et al., 2014; Rossetti et al., 2024). The relationship of the RT program to the functional regulation of the genome appears to be reciprocal, as early replication can influence the epigenome and act as a topological modulator (Chen and Buonomo, 2023; Dequeker et al., 2022; Klein et al., 2021; Stewart-Morgan et al., 2020). One of the regulators at the interplay of replication timing and the epigenome is RIF1, which is commonly found in late replicating regions in association with the PP1 phosphatase, regionally counteracting the activation of MCMs by DDK (Chen and Buonomo, 2023; Foti et al., 2016).

Studying replication of the genome in higher eukaryotes is technically challenging. The SNS-seq and INI-seq methods map replication origins at very high resolution, but their results do not match well with those obtained by other methods (Hulke et al., 2020; Tian et al., 2023). Repli-seq and OK-seq have low resolution, meaning that they define broad (50-200 kb) replication initiation zones (IZs), where several origins might be found (Petryk et al., 2018; Zhao et al., 2020). EdU-seq identifies origins at similar sites as Repli-seq and OK-seq and can also monitor fork progression. However, EdU-seq requires cell synchronisation and, to this date, only early-S initiation events have been studied by this method (Liu et al., 2021; Macheret and Halazonetis, 2019a). Unlike the origins identified in budding yeast, origins in higher eukaryotes do not appear to contain a consensus DNA sequence. Nevertheless, certain features, such as G4 structures and open chromatin marks, have been associated with origin activity (Audit et al., 2009; Brossas et al., 2020; Guilbaud et al., 2022; Jaksik et al., 2023; Long et al., 2020; Prorok et al., 2019; Smith et al., 2016).

Pluripotent stem cells constitute a powerful model to study replication dynamics, because they are highly proliferative and have a very short G1 phase (Becker et al., 2010; Roccio et al., 2013), which facilitates cell synchronisation and analysis of DNA replication. They also have a permissive chromatin environment that reflects their plasticity and pluripotency (Apostolou and Hochedlinger, 2013; Bulut-Karslioglu et al., 2018; Chen and Dent, 2013). Their permissive chromatin state is maintained by pluripotency factors, among them OCT4, SOX2, NANOG and KLF4, which act as pioneer transcription factors (Friman et al., 2019; Mayran and Drouin, 2018; Soufi et al., 2012). In this study, we applied pharmacological and genetic approaches to show that pluripotency factors, and particularly OCT4, affect the replication timing program by facilitating firing of DNA replication origins located in late S phase replicating genomic domains. Thus, we propose that pluripotency factors affect not only the transcriptional program of mESCs, but also the efficiency of the late RT program, which is further regulated by CDC7, CDK1 and ATR.

## RESULTS

### Replication timing landscape of mESCs

As a first approach to study the kinetics of genome duplication of mESCs we employed flow cytometry to determine the length of S phase. MEFs and bone-marrow mesenchymal stem cells (mMSCs) were also examined. For all three cell types, unsynchronized cells were pulsed with EdU for 30 min and their progression through the cell cycle was followed over time by flow cytometry. The EdU-positive cells reached G2 within 8-9 hours (h) in all three cell types (Supplementary Fig. 1a). Consistent with mESCs having very short G1 phases (Coronado et al., 2013; Li et al., 2012; Pauklin and Vallier, 2013; Roccio et al., 2013; Sela et al., 2012), the EdU-positive mESCs initiated a second round of DNA replication, ahead of the mMSCs and MEFs (Supplementary Fig. 1a; see vertical arrows at 9 and 12 h after the EdU pulse). In the same experiment, we also monitored progression through S phase of the EdU-negative cells. These cells entered S phase after the EdU pulse was administered and their progression through S phase, like that of the EdU-positive cells, could be monitored by the increase in their genomic DNA content (Supplementary Fig. 1a, see oblique arrows at 8 h). The profiles of the EdU-negative cells were also consistent with the length of S phase being similar in mESCs, mMSCs and MEFs.

Next, we employed high-throughput sequencing methods to monitor RT. First, we utilized a variant of Repli-seq, by labelling unsynchronized cells with EdU for 30 min and fractionating the cells based on genomic DNA content on a flow sorter (Supplementary Fig. 1b). Nascent DNA from these cells was conjugated to biotin, isolated and sequenced. From the Repli-seq profiles of pluripotent mESCs, MEFs and mMSCs, we determined the early, mid and late S phase RT domains. These domains varied somewhat across the three cell types yet overlapped significantly (Supplementary Fig. 1c) and involved about similar fractions of the genome in all cell types (Supplementary Fig. 1d).

Repli-seq experiments extrapolate time from genomic content. We, therefore, performed an EdU-seq time-course experiment, as a second approach to study RT. mESCs, synchronized in mitosis with nocodazole, were isolated by mitotic shake-off and then released in the cell cycle by washing nocodazole away. At various time points after exit from mitosis, EdU was added to the media and 30 min later the cells were harvested and processed for sequencing of their nascent DNA (Fig. 1a, b). MEFs and mMSCs could not be synchronized by mitotic shake-off, limiting the EdU-seq time-course analysis to the mESCs. As early as 2 h after mitotic exit, sharp peaks of EdU incorporation were evident, consistent with the earliest wave of origin firing and hence confirming the short G1 phase of mESCs. From 4 to 12 h after mitotic exit, the EdU incorporation profiles showed replication progressing from the early to the mid and the late S phase replication domains (Fig. 1c). Comparison of the Repli-seq (Supplementary Fig. 1c) and EdU-seq (Fig. 1c) profiles of mESCs revealed a high degree of similarity (Supplementary Fig. 1e).

**Figure 1.**
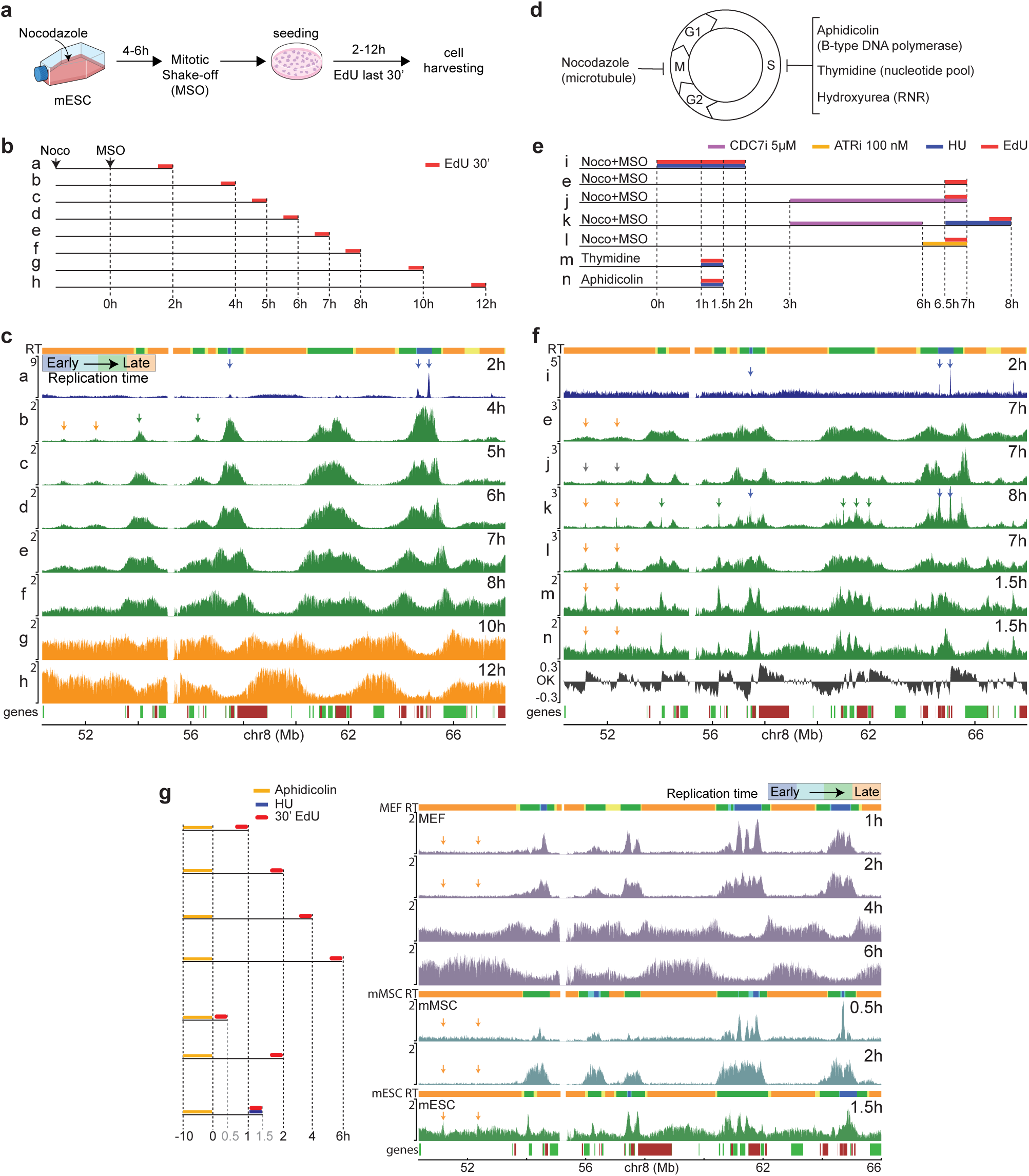
Identification of early, mid and late initiation zones (IZs). (**a**) Mitotic shake-off (MSO) scheme. (**b**) Experimental outline of the EdU-seq time-course experiment (tracks a-h, 2-12 h after MSO). EdU was added 30 minutes before harvesting the cells. (**c**) EdU-seq profiles at the indicated time points after mitotic exit. The EdU-incorporation profiles are colored blue, green or orange depending on whether the cells were in early, mid or late S phase, respectively, at the indicated time point after mitotic exit. Select IZs mapping to early, mid and late replicating regions are indicated by blue, green and orange arrows, respectively. Top: RT domains, as determined by Repli-seq (early, mid and late domains are colored blue, green and orange, respectively). Bottom: gene annotation (forward and reverse direction of transcription are marked by green and red boxes, respectively). (**d**) Scheme showing the agents used to synchronize mESCs in mitosis or at the G1/S boundary. (**e**) Experimental outline used to identify IZs at mid and late RT domains. (**f**) EdU-seq profiles in cells released into S phase from the G1/S boundary. The EdU-seq profile in tracks e is reproduced from panel (c). Tracks i-n correspond to the experiments outlined in panel (e). Track OK shows origin-mapping profiles identified by the OK-seq method (Petryk et al., 2018). Track e is the same as in panels (b and c). Select IZs are indicated by vertical blue, green and orange arrows, respectively. RT domains and gene annotation are as in panel (c). (**g**) EdU-seq profiles of MEFs, mMSCs and mESCs synchronized with aphidicolin, released and collected at different time points. Orange arrows show the positions of IZs in late replicating domains present only in mESCs (also shown in panels c and f). RT domains for each cell type are shown above their respective EdU-seq profiles.

### Late S phase IZs fire early in mESCs

The EdU-seq profiles of the mESCs revealed origin firing at multiple time points after mitotic exit (Fig. 1c). We expected that the IZs located within the mid and late RT domains would fire in mid and late S phase, respectively. However, at 4 h after mitotic exit, i.e. 2 h into S phase (corresponding to fraction S1 in the Repli-seq experiment; Supplementary Fig. 1e), we observed firing of IZs mapping within both the mid and late RT domains (Fig. 1c; green and orange arrows, respectively).

To determine whether the IZs that fired within the late RT domains in the above experiment were bona fide IZs, we examined mESCs synchronized under different conditions (Fig. 1d, e). First, we applied a CDC7 inhibitor (CDC7i) (Sotiriou et al., 2016) 3 h after mitotic exit, when the early origins had already fired, and examined nascent DNA synthesis by EdU-seq 4 h later. The CDC7i prevented the firing of the mid and late IZs, as should be the case for bona fide IZs (Fig. 1e, f: tracks e versus j). In a follow-up experiment, we released mESCs from the CDC7i block described above and monitored origin firing 2 h later. To obtain sharp EdU incorporation peaks and thus increase the resolution, we treated the cells with hydroxyurea (HU) during the release period (Fig. 1e: track k). Under these conditions, we observed the same IZs, mapping within mid and late RT domains, as the ones identified previously in untreated cells 4 h after mitotic exit (Fig. 1f: track k). The same IZs were also observed in cells synchronized by mitotic shake-off and treated with an inhibitor of the ATR checkpoint kinase to induce a burst in origin firing (Fig. 1e, f: track l). From these results we concluded that the observed peaks in the mid and late RT domains were genuine IZs. To facilitate the further study of the IZs, we classified them as early, mid or late (blue, green and yellow arrows, respectively, in Fig. 1c, f) according to the RT domain (Supplementary Fig. 1d and top bar in Fig. 1c, f) to which they mapped (see methods and Supplementary Fig. 2a, b). In total, we identified 5,179, 949 and 330 IZs mapping to early, mid and late RT domains, respectively. We next explored the dynamics of EdU incorporation associated with all the well-defined IZs. All of them contributed to the DNA replication profiles in accordance with the RT domain to which they belonged (Supplementary Fig. 2c).

Having established that well-defined IZs within late RT domains had fired in early-mid S phase in mESCs, we examined if we could obtain similar results with other cell synchronization methods. Thymidine and aphidicolin were used to induce cell cycle arrest at the G1/S boundary. Within 1.5 h of release from a thymidine or aphidicolin-induced arrest, we observed origin firing within the late RT domains (Fig. 1e, f: tracks m, n). Importantly, the IZs identified by all the experiments described above align well across experiments and also with the IZs identified in mESCs by the OK-seq method (Fig. 1f; compare track OK (Petryk et al., 2018) to tracks i-n).

Next, we investigated if the firing of late RT IZs in early-mid S phase was a specific feature of mESCs. We were unable to synchronize MEFs and mMSCs by mitotic shake-off; however, we could synchronize these cells with aphidicolin. Neither the MEFs, nor the mMSCs fired origins within late RT domains 2 h after release from an aphidicolin block (Fig. 1g, orange arrows). Instead, 4 to 6 h after release from the aphidicolin block, the MEFs exhibited broad regions of EdU incorporation in the late RT domains, suggesting different patterns of origin firing from cell to cell in these domains. Thus, firing of well-defined late RT IZs in early-mid S phase was a specific feature of mESCs.

### The RT program of mESCs is not affected by nucleotide levels

Increased levels of deoxynucleotide triphosphates (dNTPs) stimulate premature firing of late origins in yeast (Forey et al., 2020), prompting us to examine if firing of late IZs in early-mid S phase in mESCs was linked to dNTP levels. First, we supplemented the tissue culture media with nucleosides (Embryomax). This intervention did not change the RT program, as determined by both Repli-seq and EdU-seq (Supplementary Fig. 3).

In a reciprocal experiment, mESCs were treated with low doses of HU (100 μM) to impede dNTP production. The HU treatment slowed progression through S phase (Fig. 2a and Supplementary Fig. 4a-c) and lowered fork speed rates (Fig. 2b, c). However, the replication profiles, determined at various times after mitotic exit, showed no change in the RT program, other than the overall delay in S phase progression. As an example, the EdU-seq profile 6 h after mitotic exit in the untreated cells was identical to the 10 h profile of the HU-treated mESCs (Fig. 2a and Supplementary Fig. 4d-g).

**Figure 2.**
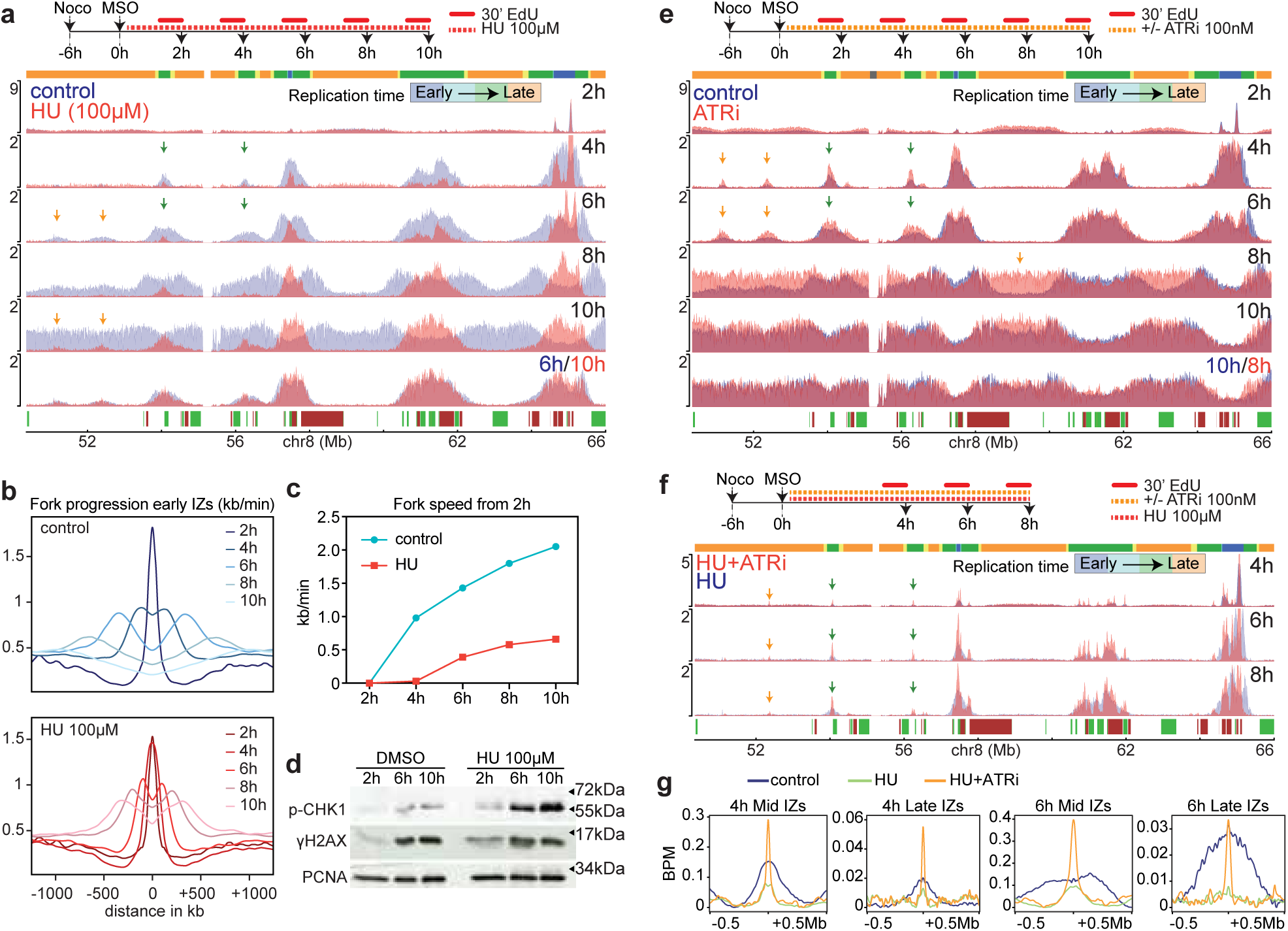
Hydroxyurea and ATR effects on firing of late origins in mESCs. (**a**) Overlay of EdU-seq plots of control (blue) and HU-treated (red) mESCs at different time points after MSO. Note that the mid and late IZs present at 6 h in control cells (green and orange arrows) could only be spotted at 8 and 10 h in the HU profiles. Above, experimental design of the experiment; EdU was added for the last 30 minutes before harvesting the cells. RT domains and annotated genes are as in Fig. 1c. (**b**) Average EdU-seq profiles around a set of isolated early origins (n=380) in control (top) and HU-treated (bottom) cells at the indicated time points after MSO. (**c**) Fork speeds calculated from the average EdU-seq profiles shown in (b). The profile at 2 h served as the starting point from which fork speeds were calculated for the later time points. (**d**) Western-blot of mESCs at 2, 6 and 10 h after MSO. The cells were treated with DMSO or HU. Molecular weight markers are shown on the right of the blot. (**e**) Overlay of EdU-seq profiles of control (blue) and ATRi-treated (red) mESCs. Green and orange arrows at 4h and 6h denote well-defined mid and late IZs. The orange arrow at 8h shows earlier replication of a late domain in ATRi-treated cells. RT domains and gene annotation are indicated as in Fig. 1c. The experimental scheme is shown above the EdU-seq profiles. (**f**) Overlay of EdU-seq profiles of mESCs treated with HU (blue) or HU plus ATRi (red) at 4, 6 and 8 h after MSO. Mid and late IZs that fired only in the HU+ATRi-treated cells are indicated by green and orange arrows, respectively. RT domains and gene annotation are as in Fig. 1c. The experimental scheme is shown above the EdU-seq profiles. (**g**) Average (BPM) EdU-seq profiles around mid and late IZs at 4 and 6 h after MSO in control, HU and HU plus ATRi-treated mESCs.

### Interplay between ATR, CDC7 and CDK1 in regulating late origin firing

The DNA replication checkpoint has been reported to control the timing of late origin firing via pathways that involve CDC7, RIF1 and CDK1. The main effector of the DNA replication checkpoint is the kinase ATR (Saldivar et al., 2017). Upon entry into S phase, exhaustion of dNTPs, as a result of DNA synthesis, activates ATR, which triggers dNTP synthesis (Forey et al., 2020; Le et al., 2017; Sugitani et al., 2022). In parallel, ATR suppresses firing of more origins until appropriate dNTP levels are reached. The molecular mechanism involves phosphorylation of DBF4, the regulatory subunit of CDC7 (Heffernan et al., 2007; Zegerman and Diffley, 2010), and activation of RIF1, which associates with Protein Phosphatase 1 at late origins to antagonize CDC7 activity (Moiseeva et al., 2019; Rainey et al., 2020; Seller and O’Farrell, 2018; Simoneau and Zou, 2021).

We wondered whether mESCs lacked a functional DNA replication checkpoint, as this could explain the firing of late IZs in early-mid S phase. As a first step to answer this question, we determined whether the DNA replication checkpoint is activated in unperturbed mESCs during S phase by monitoring phosphorylation of histone H2AX and the kinase CHK1, both of which are ATR substrates (Liu et al., 2000; Saldivar et al., 2017). Consistent with previous reports (Ahuja et al., 2016), at 6 and 10 h post-mitotic exit, we observed increased levels of phosphorylated H2AX (γH2AX) and CHK1 (p-CHK1) in both untreated mESCs and mESCs treated with low doses of HU (Fig. 2d). In subsequent experiments, inhibition of ATR by a specific chemical inhibitor (ATRi; BAY-1895344) stimulated firing of mid and late IZs, both in the presence and absence of low doses of HU (Fig. 2e-g; green and orange arrows for mid and late IZs, respectively). We conclude that the DNA replication checkpoint is not defective in mESCs and that it curtails firing of mid and late IZs, making ATR a putative rheostat for the transition into the late replication program.

In addition to the DNA replication checkpoint, firing of late origins is regulated by the CDC7-RIF1 axis and by CDK1 (Moiseeva et al., 2019; Rainey et al., 2020; Simoneau and Zou, 2021). To study the interplay between ATR, CDC7 and CDK1, we synchronized mESCs by mitotic shake-off and 3 h later treated them with TAK-931, a highly specific CDC7 inhibitor. EdU-seq profiles from cells harvested 4 and 7 h after mitotic exit revealed that the CDC7 inhibitor suppressed firing of origins at the mid and late RT domains (Fig. 3a, b). In agreement with the DNA replication checkpoint antagonizing CDC7, inhibition of ATR rescued firing of these origins in mESCs treated with TAK-931 (Fig. 3a, b). Moreover, the rescue of origin firing was suppressed by a CDK1 inhibitor (Vassilev et al., 2006) (Fig. 3a, b), consistent with reports that ATR inhibitors activate CDK1, which in turn induces origin firing by inhibiting RIF1 and by phosphorylating the MCM helicase (Fig. 3c) (Komamura-Kohno et al., 2006; Lin et al., 2008; Masai et al., 2000; Moiseeva et al., 2019; Rainey et al., 2020; Seller and O’Farrell, 2018; Simoneau and Zou, 2021; Suski et al., 2022).

**Figure 3.**
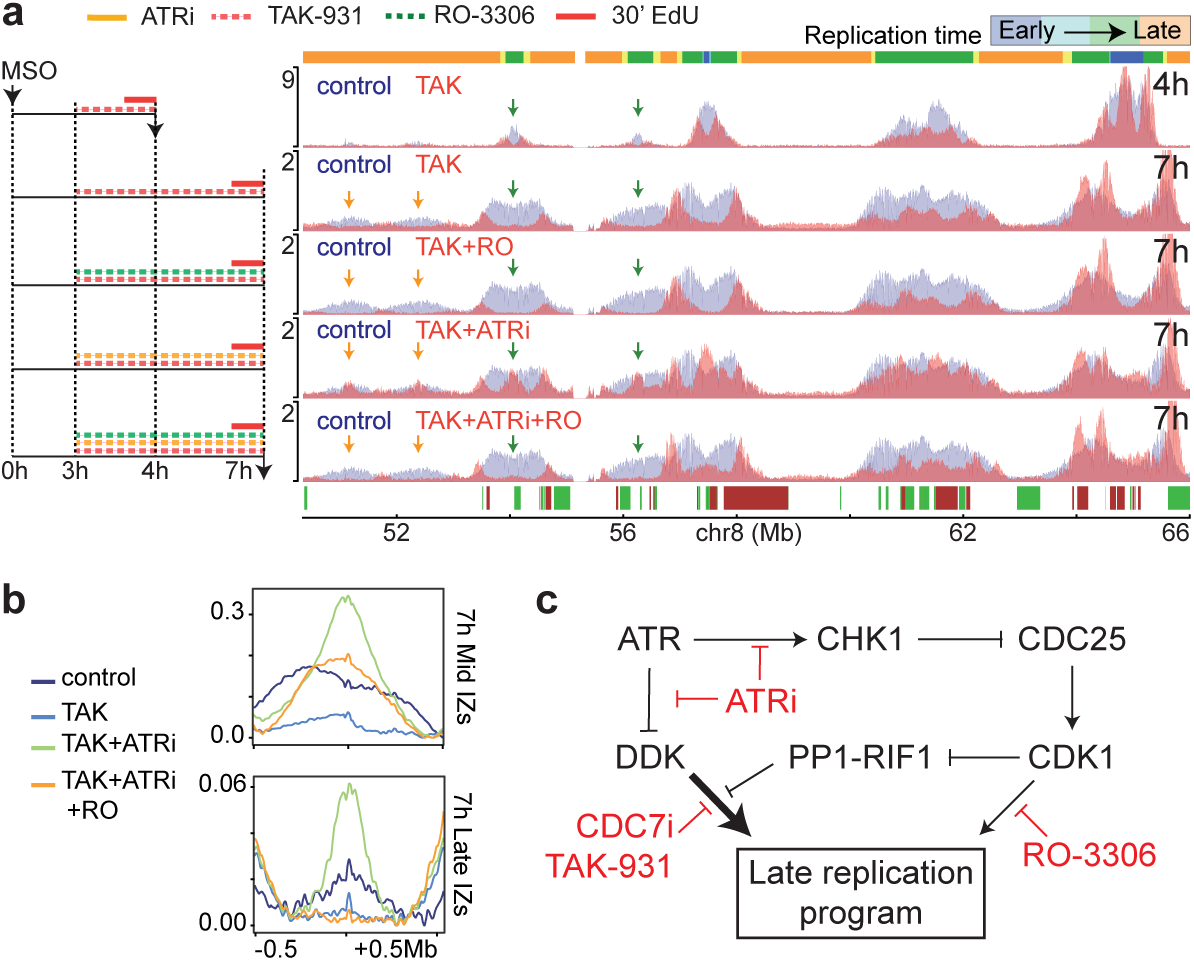
DDK and CDK1 induce firing of late IZs. (**a**) EdU-seq profiles of mESCs treated with CDC7 inhibitor (TAK-931, TAK), ATR inhibitor (ATRi) and/or CDK1 inhibitor (RO-3306, RO). Left, experimental outlines; right, corresponding EdU-seq profiles. Green and orange vertical arrows correspond to select mid and late IZs, respectively. RT domains and gene annotation are indicated as in Fig. 1c. (**b**) Average EdU-seq profiles around mid and late IZs 7h after MSO for control mESCs and mESCs treated with the indicated inhibitors. (**c**) Molecular pathways by which ATR regulates late origin firing. DDK, CDC7 and DBF4 complex; PP1, protein phosphatase 1.

### A high fraction of the genome is transcriptionally active in mESCs

Most transcriptionally active genomic domains are replicated in early S phase (Goldman et al., 1984; Hiratani et al., 2008; Marchal et al., 2019b; Schwaiger et al., 2009; White et al., 2004). Therefore, the firing of late IZs in early-mid S phase in mESCs might be related to a higher fraction of the genome being transcriptionally active in mESCs, as compared to differentiated cells. To address this possibility, we monitored nascent transcription in mESCs and MEFs. The uridine analogue 5-ethynyl-uridine (EU) was added to cultures of asynchronous cells for 30 min; the cells were then sorted by their genomic content into six fractions, as done for Repli-seq (Fig. 4a), and nascent transcription was determined genome-wide for each fraction by EU-seq.

**Figure 4.**
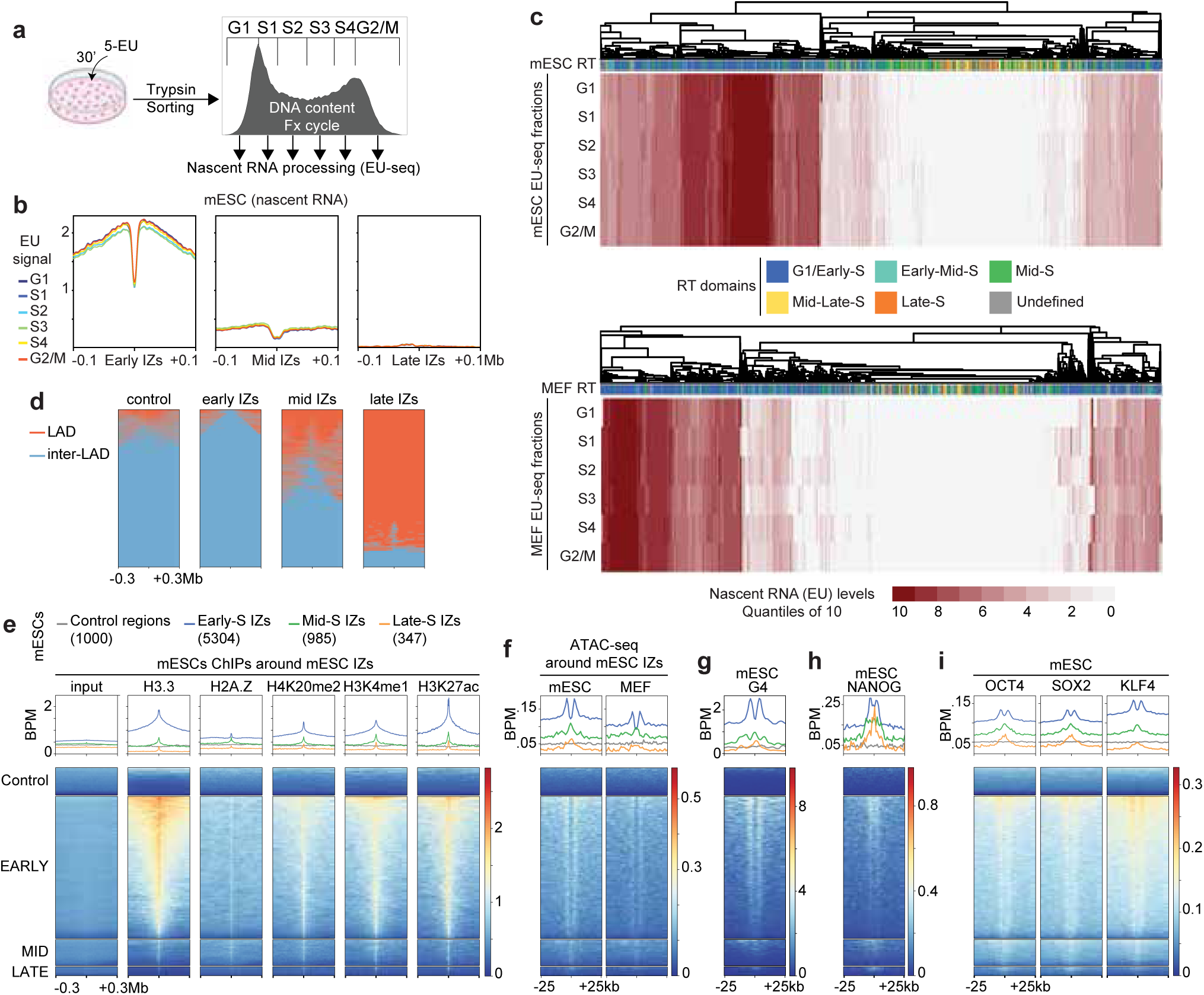
Transcription patterns and open chromatin marks at IZs. (**a**) Experimental outline of EU-seq experiments with sorted cells. EU was added for 30 minutes to non-synchronized cells (mESCs or MEFs); the cells were then sorted in six fractions according to DNA content (compare to Supplementary Fig. 1b). (**b**) Average EU-seq signal around early, mid and late IZs for each fraction of sorted mESCs. (**c**) Gene clustering according to nascent transcription along the cell cycle in mESCs (left) and MEFs (right). The genes (displayed in horizontal rows) were clustered according to their relative expression levels in the six cell cycle fractions (columns G1 to G2/M) and RT (first column of the heatmap). (**d**) Heatmaps showing lamina-associating domains (LADs) and inter-LAD regions (Peric-Hupkes et al., 2010) around early, mid and late IZs. (e) Heatmaps showing ChIP-seq signal intensities for the indicated histone marks around all early, mid and late IZs from published mESC datasets. The number of plotted early, mid and late IZs are shown in parentheses. (**f**, **g**, **h**, **i**) Heatmaps of ATAC-seq signals from mESC and MEF (f), G4 quadruplexes (e) and ChIP-seq signals of NANOG (h), OCT4, SOX2 and KLF4 (i) from published mESC datasets at early, mid and late IZs.

The genome-wide averages of EU signals around all early, mid and late IZs in mESCs revealed high levels of nascent transcription flanking the early IZs, a modest signal around the mid RT IZs and practically no signal around the late IZs (Fig. 4b). We note that there was a sharp drop of nascent transcription signal at the very center of the early and mid IZs (Fig. 4b), which can be explained by replication origins mapping to transcriptionally silent intergenic regions next to actively transcribed genes (Cayrou et al., 2011; Macheret and Halazonetis, 2018; Martin et al., 2011; Rossetti et al., 2024; Sequeira-Mendes et al., 2009).

Next, we compared the nascent transcription landscape of mESCs and MEFs. For each cell type, we clustered all the annotated transcripts according to nascent transcription levels, flow sorting fraction (based on genomic DNA content) and RT (Fig. 4c). In both mESCs and MEFs, early RT domains were enriched with mid and highly transcribed genes, whereas genes located in late replicating domains were generally not expressed (Fig. 4c). Of the early-replicating genes, more were transcriptionally active in mESCs than in MEFs; however, the late-replicating genes were not transcribed in either the mESCs or the MEFs (Fig. 4c). Thus, transcriptional activity cannot explain why firing of late S IZs differs between mESCs and MEFs.

### Epigenetic marks at IZs

Since we observed origin firing even in non-transcribed genomic domains, we next examined whether IZs were associated with features of open chromatin. To address this possibility we analysed publicly available mESCs epigenetic data (Chronis et al., 2017; Juan et al., 2016; Lyu et al., 2022; Navarro et al., 2020; Peric-Hupkes et al., 2010; Wen et al., 2020). We saw that early, mid and late IZs were distributed differently across lamina-associating domains (LADs) (Peric-Hupkes et al., 2010; Pope et al., 2014). Thus, most early and mid IZs mapped to broad inter-LAD regions; whereas, the late IZs were mostly located within LADs (Fig. 4d).

The early IZs were decorated with active histone variants (H3.3 and H2AZ) and euchromatic marks (H3K4me1 and H3K27ac) (Fig. 4e and Supplementary Fig. 5). H4K20me2, which is recognized by the BAH domain of ORC1, was also enriched at the early IZs (Fig. 4e), as were G4 quadruplexes and open chromatin regions identified by the Assay for Transposase-Accessible Chromatin (ATAC-seq) (Fig. 4f, g). The same marks that were enriched at the early IZs were also enriched at the mid and late IZs, although the signal intensity was lower (Fig. 4e, f, g and Supplementary Fig. 5). Thus, open chromatin features are present at IZs and their signal intensity correlates with RT.

### Pluripotency factors act as pioneer factors for mid and late IZs

Given the presence of open chromatin marks at IZs, we wondered whether the firing of late IZs in early-mid S phase might be driven by recruitment of mESC-specific pioneering factors. Likely candidates would be the pluripotency factors OCT4, SOX2, KLF4 and NANOG, which are expressed in mESCs, but not in differentiated cells. Implicit in this hypothesis is that the abovementioned proteins bind to the genomic sites, where the mid and late IZs are located. Analysis of publicly available datasets (Chronis et al., 2017) revealed that these pluripotency factors are enriched at early, mid and late IZs (Fig. 4h, i and Supplementary Fig. 5).

Out of the above pluripotency factors, we focused on OCT4 by studying the well-characterized ZHBTc4 stem cells, in which OCT4 levels are regulated by a tet-off system (Fig. 5a) (Niwa et al., 2000). Treatment of these cells with doxycycline for 12 or 18 h suppressed OCT4 levels (Supplementary Fig. 6a) without affecting the kinetics of entry into S phase after synchronization by MSO (Supplementary Fig. 6b). To monitor firing of mid and late IZs, the ZHBTc4 cells were treated with or without doxycycline and synchronized by MSO or with thymidine. Samples were collected at several time points after release from the cell cycle block and processed for EdU-seq. Inspection of representative genomic regions revealed delayed firing of mid and late IZs in the absence of OCT4 (Fig. 5c, d and Supplementary Fig. 6c). The affected IZs corresponded to sites where OCT4 bound to regulate chromatin accessibility, as documented from publicly available datasets (Chronis et al., 2017) (Fig. 5c, d and Supplementary Fig. 6c).

**Figure 5.**
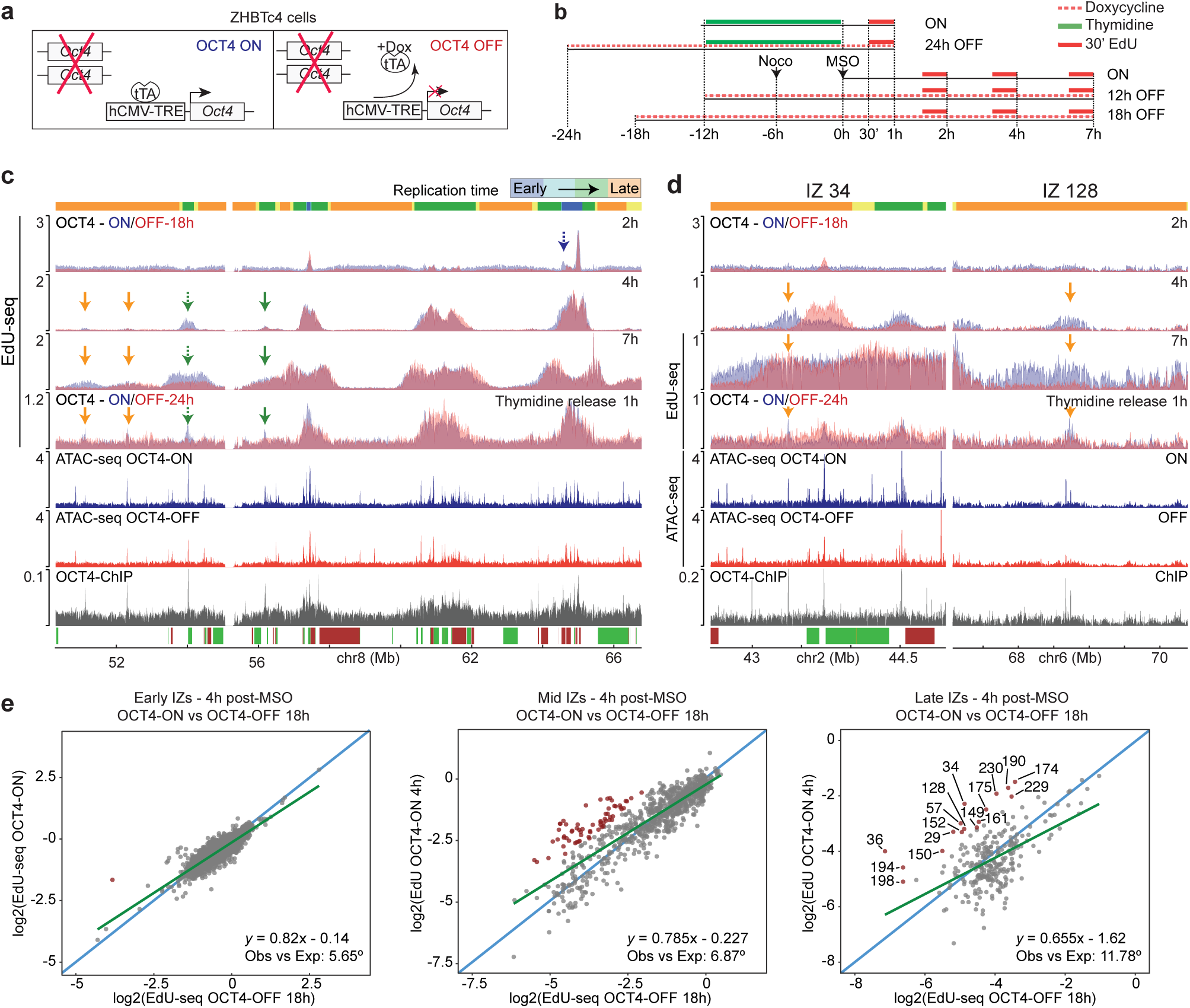
OCT4 enhances firing of select mid and late IZs in mESCs. (**a**) Scheme of the tet-off ZHBTc4 cells, in which OCT4 levels can be depleted by adding doxycycline (Dox). (**b**) Outline of EdU-seq experiments performed with ZHBTc4 cells synchronized with thymidine or by mitotic shake-off (MSO). To suppress OCT4 expression, treatment of the cells with doxycycline was initiated 24 h before release from the thymidine block or 12 or 18 h before the MSO and kept until the cells were harvested. (**c**) EdU-seq profiles of ZHBTc4 cells at the indicated times after MSO or after release from a thymidine block. Profiles with OCT4 expression turned ON or OFF (doxycycline added 18 h before MSO) are compared. Plots below the EdU-seq profiles: ATAC-seq profiles from asynchronous ZHBTc4 cells with OCT4 expression turned ON or OFF (Friman et al., 2019) and OCT4 ChIP-seq profile from asynchronous mESCs (King and Klose, 2017). Select IZs mapping to mid and late replicating regions are indicated by green and orange arrows, respectively. RT domains and gene annotation are indicated as in Fig. 1c. (**d**) EdU-seq, ATAC-seq and OCT4 ChIP-seq profiles for the genomic regions spanning IZs 34 and 128, shown as in panel c. (**e**) Scatterplots of EdU-seq signals 4 h after MSO at early, mid and late IZs of ZHBTc4 cells expressing (ON) versus not expressing (OFF) OCT4. OCT4 expression was suppressed by administering doxycycline 18 h before MSO. The EdU-seq signal for each IZ was computed as BPM in a region +/-5 kb around the centre of the IZ and transformed into log2. The diagonal blue lines indicate a hypothetical perfect correlation; the green lines show the trend of the real data; the angle between the lines was calculated to highlight the differences. All IZs with ON/OFF rates >2.5 are highlighted in red. Some late IZs are numbered for referencing.

For a genome-wide analysis of the effect of OCT4 on the initiation of replication, we generated scatter plots of the EdU-seq signal 4 h post-MSO at all early, mid and late IZs in ZHBTc4 cells treated with or without doxycycline. The absence of OCT4 had little effect on firing of the early IZs, but markedly suppressed firing at select mid and late IZs (Fig. 5e and Supplementary Fig. 6d). The duration of the treatment with doxycycline, 18 or 12 h, did not affect the results (Fig. 5e and Supplementary Fig. 6d, respectively).

Scatter plots of the ATAC-seq signal from public datasets paralleled the EdU-seq data, as they revealed decreased chromatin accessibility at late IZs following OCT4 depletion (Supplementary Fig. 6e). Considering that OCT4 was present at most of the genomic sites corresponding to mid and late IZs (as examples see the sites labelled IZ34 and IZ128 in Fig. 5d), our results suggest that OCT4 enhances chromatin accessibility at mid and late S phase IZs and in doing so increases origin firing efficiency, thus promoting their activation in early S phase.

## DISCUSSION

In this study, we investigated the replication dynamics of pluripotent cells. Remarkably, we observed that pluripotency factors stimulate the firing of origins within late replicating genomic domains, suggesting that these factors have functions that extend beyond regulating transcriptional activity.

mESCs provide a unique system to study DNA replication IZs. Because the G1 phase of these cells is only about 2 h long, mESCs enter S phase rather synchronously after mitotic exit. This allowed us to use the EdU-seq method ^61^ to monitor origin firing upon entry of the cells into S phase and at later time points, without having to treat the cells with HU. As a result, we observed origin firing within the mid and late RT genomic domains and were able to map these IZs on the genome with high resolution.

Our results shed light on general principles defining the initiation of DNA replication and the establishment of the RT program (Fig. 6). In mESCs, DNA replication was mediated by well-defined IZs. The early RT IZs were very efficient and fired shortly after entry into S phase. The firing of mid RT IZs was evident at 3 h and continued up to 8 h after mitotic exit, indicating a lower efficiency of origin firing that took several hours to complete. The late RT IZs were even less efficient with firing evident in few cells as early as 4 h and continuing beyond 8 h after mitotic exit. Our results fit into the “controlled-stochastic” model of eukaryotic DNA replication, a model supported by single-cell and single-molecule approaches, which posits that the efficiency of origin firing is a determinant of RT (Dileep and Gilbert, 2018; Gnan et al., 2022; Hu and Stillman, 2023; Raghuraman and Brewer, 2010; Rhind et al., 2010; Takahashi et al., 2019; van den Berg et al., 2024; Wang et al., 2021).

**Figure 6.**
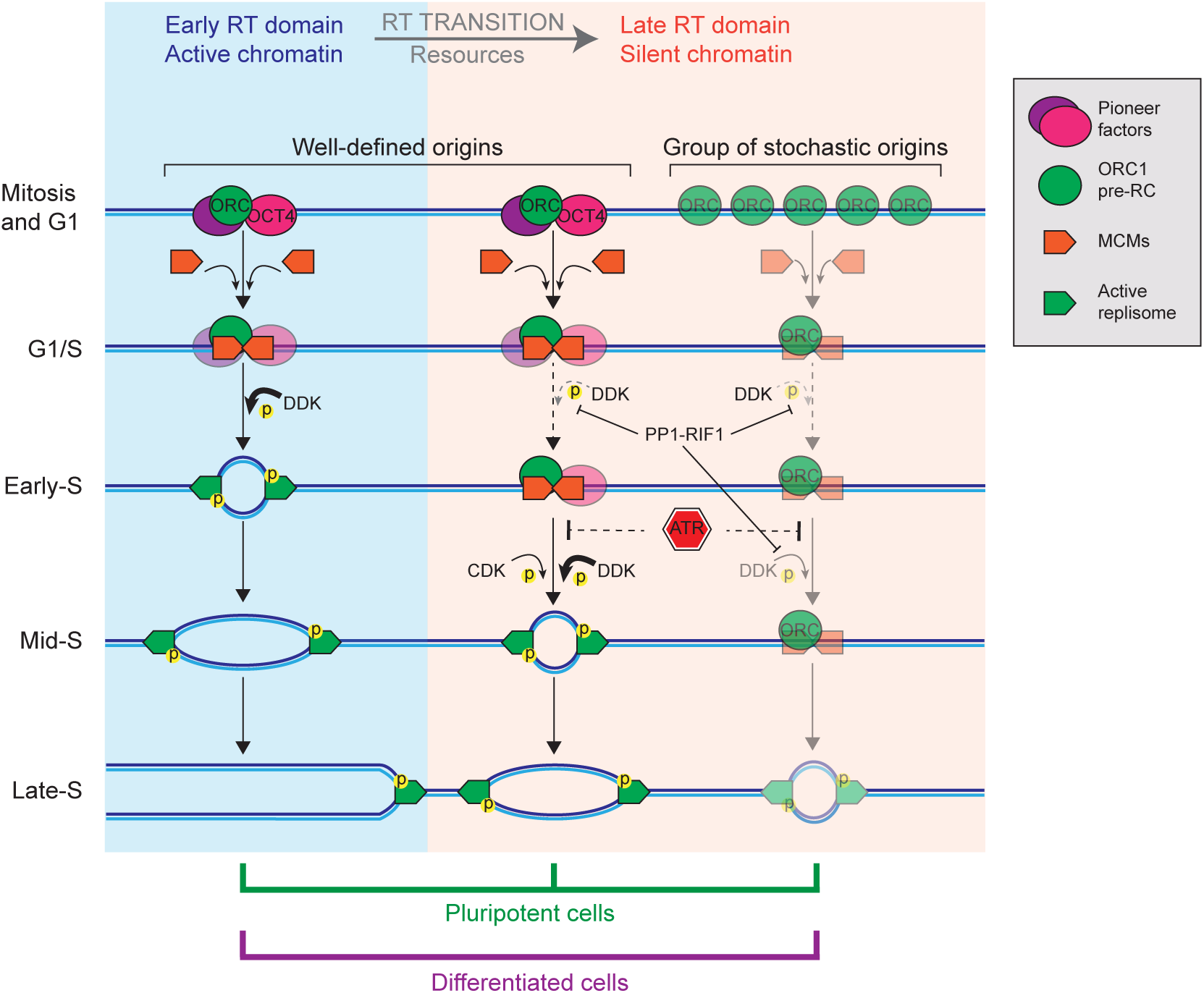
Model for origin firing and RT. Several layers control the efficiency and temporality of origin firing: locally, efficient origin firing is favoured by chromatin accessibility; globally, the transition into the late replicating program is controlled by ATR, RIF1 and the availability of resources. In mESCs, efficient, well-defined origins can be found in early, mid and late RT domains. In late RT domains, the firing of some of these origins depends on local recruitment of the pluripotency factor OCT4. In differentiated cells, efficient, well-defined origins can be found only in early and mid RT domains and replication of the late RT domains depends on low efficiency origins that cannot be mapped, when populations of cells are analysed.

We did not elucidate here what drives the higher firing efficiency of early IZs, as compared to the less efficient mid and late IZs. Nevertheless, we note that the firing efficiency correlated with chromatin accessibility, as the signal intensity of open chromatin marks correlated well with RT and, by inference, with firing efficiency. In yeast, it is documented that an open chromatin structure contributes to origin firing efficiency by increasing the probability of MCM recruitment (Das et al., 2015; Dukaj and Rhind, 2021; Foss et al., 2021).

Interestingly, not all late replicating genomic domains in mESCs were associated with a late RT IZ, as some late RT genomic domains were replicated 10-12 h after mitotic exit without well-defined IZs being evident at earlier time points. An example is the late RT domain located at the center of the representative genomic region shown throughout this study (chr8, Mb 58-60; Fig. 1c and Fig. 1f), where replication appears to initiate simultaneously across this 2 Mb region (Fig. 1c and Fig. 1f). This pattern suggests the presence of multiple low-efficiency origins throughout late domains (Fig. 6). Under one scenario all these origins could fire simultaneously in every cell or, more likely, a few of these origins would fire in each cell in a stochastic manner (Fig. 6). Since EdU-seq monitors origin firing in a population of cells, we cannot discriminate between these possibilities.

The existence of two ways by which late RT domains are replicated in mESCs is supported by data obtained by OK-seq and single-cell methods (Gnan et al., 2022; Petryk et al., 2016; Takahashi et al., 2019; van den Berg et al., 2024; Wang et al., 2021). The late RT domains with well-defined IZs by EdU-seq match OK-seq data showing origins at the same genomic positions (for example, Fig. 1f; chr 8, Mb 51-53), whereas no origins were evident by OK-seq in the late RT domains that lacked well-defined IZs by EdU-seq (for example, Fig. 1f; chr 8, Mb 58-60).

Unlike mESCs, the late RT domains in MEFs appeared to be replicated exclusively by low-efficiency stochastic origins (Fig. 1g). We attribute the presence of well-defined late IZs in mESCs to pioneer factors, such as OCT4, that enable specific genomic sites to serve as IZs, presumably by locally opening the chromatin structure. Such a hypothesis is consistent with previous studies showing that transcription factors can stimulate origin activity in diverse organisms ranging from viruses to yeast and metazoans (Cadoret et al., 2008; Dellino et al., 2013; DePamphilis, 1993; Knott et al., 2012; Li, 1999; Marahrens and Stillman, 1992; Ostrow et al., 2017; Sima et al., 2019; Turner et al., 2023). Moreover, the acute depletion of the pluripotency factor OCT4 in mESCs led to decreased chromatin accessibility (Friman et al., 2019; Mayran and Drouin, 2018), concurrent to decreased origin firing efficiency at the well-defined late IZs (Fig. 5).

Beyond the local determinants, origin firing in mESCs was regulated by kinases that regulate cell cycle progression, including CDC7 and CDK1, and kinases that signal the presence of DNA replication stress, notably ATR. Inhibition of ATR increased origin efficiency, even when CDC7 was inhibited; under these conditions, origin firing was dependent on CDK1 activity. Our results are, thus, consistent with molecular pathways already described in yeast and mammalian cells that link CDC7, CDK1 and ATR activities (Ahuja et al., 2016; Heffernan et al., 2007; Katsuno et al., 2009; Komamura-Kohno et al., 2006; Lin et al., 2008; Liu et al., 2019; Masai et al., 2000; Michowski et al., 2020; Moiseeva et al., 2019; Rainey et al., 2020; Saldivar et al., 2017; Seller and O’Farrell, 2018; Simoneau and Zou, 2021; Suski et al., 2022; Wang et al., 2016).

In conclusion, our work suggests that RT is determined by the firing efficiency of well-defined origins and by global regulators, including the DNA replication checkpoint and cell cycle-regulated kinases. In mESCs, we have documented the presence of well-defined IZs, even in late replicating genomic domains. The presence of these efficient IZs is dependent on expression of pluripotency factors, which act as pioneering factors to open chromatin structure locally. The functional significance of these well-defined IZs in late RT domains in mESCs remains to be determined.

## MATERIAL AND METHODS

### Cell culture

Mouse embryonic stem cells (mESCs) from 129/SVEV strain were purchased from Merck (CMTi-1, passage 11). They were thawed, amplified as per manufacturer instructions for three passages, aliquoted and frozen. This stock of cells was routinely used on gelatin plates (or flasks) for no more than 32 passages. For all experiments, mESCs were grown in DMEM+Glutamax (Gibco, 61965-026), 10% ES-FBS (Gibco 16141-079), MEM NEAA (Gibco 11140-035), sodium pyruvate (Gibco, 11360-039), 1mM beta-mercaptoethanol (Gibco, 31350-010), penicillin/streptomycin (P/S, Gibco, 15140-122), 100ng/mL LIF (stocked at 1-2mg/mL; produced at the Protein Production and Purification Platform of the EPFL, Lausanne, Switzerland), 3µM GSK3 inhibitor (Sigma 361559) and MEK inhibitor (PD0325901, Selleckchem). ZHBTc4 (Niwa et al., 2000) are tet-off mouse pluripotent cell lines which were obtained from David Suter (EPFL, Switzerland). They were also grown on gelatin plates using the same media as mESCs. To deplete OCT4 (ZHBTc4), doxycycline was added to the growing media at the specified times (see figures and text).

Mouse embryonic fibroblasts (MEFs) were obtained from eviscerated torsos of E12-14 mouse embryos. Tissue from three embryos was minced with a scalpel on a plate with 2-3 mL of trypsin. The minced tissue was then placed in a 50 mL falcon tube containing 10 mL of trypsin for 10-15 minutes at 37°C, followed by strong pipetting and another 10-15 minutes incubation at 37°C. 20 mL of complete DMEM (DMEM, 10% FBS, P/S, L-glutamine) were added to the cell suspension, which was subsequently pipetted again and let to sit for 5 minutes. The supernatant was then recovered and centrifuged 5 minutes at 300G. The cell pellet was resuspended vigorously and plated on a 10 cm plate. This was considered passage 0 and from here MEFs were passaged, amplified, aliquoted and frozen. They were routinely cultured in complete DMEM and used for all experiments at passage three or four, except for the EU and γH2AX FACs experiments for which they were used at passages three and ten. Bone-marrow mouse mesenchymal stem cells (mMSCs) were a kind gift from Francesc Ventura (University of Barcelona, Spain). They had been obtained from the bone marrow of 8-10 week-old mice and were grown in complete DMEM with 15% FBS.

All cells were routinely checked for mycoplasma contamination.

### Synchronisation methods

Mitotic shake-off was conducted on mESCs and ZHBTc4 as follows. Three to four million cells were seeded in T225 flasks using 40 mL of media. Two days later, 100ng/mL nocodazole was added to the media for 4 to 6 hours. Media was then removed, and 15 mL of PBS were carefully added to the cells. Round cells were lifted by tapping the sides of the flasks and visually inspected. Depending on the conditions of the experiment, cell suspensions from different flasks were strained through a 40 µm mesh and pooled into 50 mL tubes, centrifuged at room temperature (400G, 5 minutes), washed once with 15 mL of PBS and centrifuged again. Pelleted cells were then carefully resuspended and plated. Commonly, the cells lifted from two T225 flasks would be seeded on one 10 cm plate (for EdU-seq and EU-seq) or on one 6 or 12 multi-well plate (for cell cycle analysis).

For thymidine or aphidicolin synchronisation, pluripotent cells (0.5-1×10^6^ mESCs or ZHBTc4) or MEFs (1.5-2×10^6^ cells) were seeded on 10 cm plates. 24 to 48 hours later, 2mM thymidine or 4µM aphidicolin was added to the media for 10 to 12 hours. Cells were washed three to four times with PBS and released into fresh media.

### Flow cytometry and cell cycle analysis

All pluripotent cell lines used (mESCs and ZHBTc4) were synchronised by mitotic shake-off, following, as needed, a doxycycline regime for the tet-off cell line (see figures and the results section for details). Cells lifted from two T225 flasks were plated on a gelatin 12-well multi-well plate. Cells were collected with trypsin at the indicated time points, with 50 µM EdU added during the last 30 minutes. After centrifugation, cell pellets were resuspended in PBS and fixed in methanol (final concentration 90%) and stored at -20°C until further processing. The samples were centrifuged at 400G for 10 minutes at 4°C. The pellets were resuspended in 0.2% triton/PBS, transferred to a 96-well V-bottom plate and incubated at room temperature for 20 minutes. The plate was then centrifuged, supernatants removed by tilting and the sample wells washed; followed by another centrifugation, and supernatants removal. A click-IT cocktail mix was prepared containing, in one ml of final volume, 855 µL of 100mM Tris-HCl pH 8, 100 µL of 1M sodium ascorbate, 40 µL of 100mM CuSO4 and 1 µL of Alexa 647 Azide (Invitrogen A10277). Each sample/well was resuspended with 100-150 µL of the click-IT cocktail mix and incubated protected from light at room temperature for 30 minutes. The plate was then centrifuged, supernatants discarded, and pellets washed in PBS. After another centrifugation, pellets were resuspended in a solution containing 150 µL of PBS, 0.5 µg RNase (Roche, 11119915001) and 1µg/mL of propidium iodide and incubated for at least 2 h at room temperature or at 37°C. The samples were analysed in a Beckman Coulter Cytoflex device.

### Western Blot

Samples were collected with trypsin, pelleted and kept at -80°C. They were resuspended in 100 µL of lysis buffer containing 10 mM Tris-HCl pH 7.5, 50 mM NaCl, 1 mM EDTA, 1 mM DTT, 10% glycerol, 10 mM Na-molybdate, protease inhibitors (EDTA free, Pierce, A32965) and phosphatase inhibitors (Thermo, 78420) and sonicated in a Diagenode Bioruptor waterbath in two cycles of 10 minutes (30 seconds on, 30 seconds off). The soluble fraction was recovered by centrifugation (10 minutes, 12,000 G at 4°C) and protein concentration was measured by the Bradford method (Biorad, 500-0006). Samples were then boiled in SDS Novex Buffer (Thermo LC2676) at 95°C for 5 minutes. Between 20-40 µg of protein sample were loaded on each lane on 10-15% SDS-PAGE home-made gels. The wet transfer to nitrocellulose membranes (0.45 µm Porablot NCP-Macherey Nagel, 741280) took 90 minutes at a constant 250A current in VWR tanks in transfer buffer (20% methanol, Tris, glycine). A control Ponceau staining was performed for 2-3 minutes and membranes were then subsequently washed in a big volume of deionized water. Membranes were blocked for one hour with 0.5% BSA (Roche, 10735086001) in TBST buffer. Primary antibodies were incubated overnight at 4°C on a shaking platform. Membranes were washed three times in TBST and incubated in secondary antibodies-HRP (Pierce Goat anti mouse HRP cat. no. 31430 or goat anti-rabbit HRP, 31460) diluted 1:10000 in 2.5% BSA/TBST) for one hour on a shaking platform. After four TBST washes, membranes were revealed using ECL (Advantsa WITEC WesternBright, K12045-D20) and imaged in Amersham or Licor devices.

### EdU-seq

Cells were treated with 40µM of EdU for 30 minutes before collection by trypsin. EdU-seq was performed as in (Macheret and Halazonetis, 2019b) but with certain modifications. Methanol-fixed cells were pelleted, resuspended in 0.2% Triton and incubated for 20 minutes, followed by another centrifugation and one PBS wash. Cells were pelleted again, resuspended in 300-500 µL of click-IT solution and incubated for 30 minutes. Afterwards, cells were pelleted and washed, followed by lysis in 300 µL of buffer for a minimum of 3 hours at 55°C. Phenol/chloroform extraction and chloroform purification were sequentially carried in the same phase-lock tube. DNA was recovered by precipitation with NaCl and ethanol, quantified by Qubit broad range kit and kept frozen at -20°C until later use or immediately processed. Between 1-5 µg of total DNA were diluted in 105 µL of water and sonicated in a Diagenode Pico Bioruptor device for 7 cycles (15 seconds ON, 90 seconds OFF). DNA was then cleaned as specified in the abovementioned protocol using MyOne Streptavidin C1 Dynabeads and posteriorly eluted in 55 µL of 10mM Tris pH 8 and 1.1 µL 2-mercaptoethanol. DNA library preparation and sequencing were performed at the Genomics platform of the University of Geneva.

### Repli-seq

Cells were seeded in T225 flasks and grown for 48 hours after which EdU (40 µM) was added for 30 minutes. At least 20 million cells were harvested per sample. The cells were collected by trypsinization, pelleted, resuspended in PBS, fixed in methanol added drop-wise (final concentration 90%) and stored at -20°C until further processing. Cells were centrifuged at 4°C, 400G for 10 minutes, washed with 2-5 mL PBS, pelleted again and permeabilized by resuspending in 5 mL 2% triton/PBS for 30 minutes. Cells were pelleted, washed in PBS and passed into a 5 mL low bind tube. They were centrifuged, resuspended in 1-5 mL of Biotin-Azide Reaction cocktail (as in EdU-seq) and incubated for 30 minutes at room temperature. Cells were then centrifuged, washed once in PBS and resuspended in PBS containing 500µg/mL propidium iodide and 500-100U of RNaseA (Thermo EN0531). The sample was incubated for 30 minutes at 37°C and kept at 4°C until the day after. The cells were then sorted in six fractions according to the cell cycle PI profile on a MOFLO Astrios EQ (Beckman Coulter) at the Cytometry platform of the University of Geneva. At least one million cells per fraction were sorted directly into low bind centrifuge tubes, which were later centrifuged and processed as in the EdU-seq protocol.

### EU-seq

For FACS sorted EU-seq, one million mESCs or 3 million MEFs were seeded on 10 cm plates; the following day, EU (5-ethynil-uridine, Jena Biosciences, Cat. No. CLK-N002-10) was added at a concentration of 0.5 mM 30 minutes before collecting the cells by trypsinization and fixing them in methanol. The cells were stored at -20°C overnight or longer. The fixed cells were pelleted (500G, 10 minutes at 4°C), washed in cold PBS and pelleted again. The samples were then resuspended in fresh PBS, stained with FxCycle Violet Stain (F10347), fractionated as in the Repli-seq experiments, pelleted (600 G, 15 minutes, 4°C) and immediately resuspended in 600 µL Trizol. The samples were then processed as in (Macheret and Halazonetis, 2018), but with certain modifications. Briefly, total RNA was extracted through chloroform purification and ethanol precipitation. 10-15 µg of EU-labelled RNA was processed with the Click-iT Nascent RNA Capture Kit (Invitrogen, Cat. No. C-10365), except that the biotin-azide from the kit was substituted with cleavable biotin-azide (Azide-PEG(3+ 3)-S-S-biotin) (Jena Biosciences, CLK-A2112-10). The click-IT reaction took place on a volume of 50 µL for 30 minutes at room temperature. RNA samples were then precipitated by adding 350 µL of water, 16 µL of 5M NaCl, 1 µL of glycogen and 832 µL of cold ethanol, incubating the mix at -70°C and centrifuging for 30-60 minutes at 4°C, 16000 G. The RNA pellet was washed twice in a cold solution of 75% ethanol/water, centrifuged and the pellet resuspended in 400 µL of water. Biotinylated RNA was captured using 200 µL of Dynabeads MyOne streptavidin C1 (Invitrogen, 65001) per sample. The beads had been previously washed thrice in buffer W1x (5 mM Tris-HCl pH 7.5, 0.5 mM EDTA, 1 M NaCl, 0.5% Tween-20), twice in Solution A (0.1M NaOH, 0.05M NaCl in water) and twice in Solution B (0.1M NaCl in water). The volume of solutions used for washing the beads was calculated as 200 µL per sample and scaled up depending on the total number of samples. After the last wash, the beads were removed from the magnet and resuspended in buffer W2x (same as W1x but two times concentrated) in two times the original volume (400 µL per sample). 400 µL of this washed bead mix were added to each biotinylated RNA sample and incubated 30 minutes at room temperature on a rotating wheel. Afterwards, the bead-RNA samples were washed three times with 800 µL of buffer W1x. Each Biotinylated-RNA sample was eluted by resuspending the beads in a solution containing 55 µL of 10mM Tris pH 8 and 1.1 µL of 2-mercaptoethanol and incubating them for one hour at room temperature. RNA libraries were generated without the ribodepletion step and sequenced at the Genomics platform of the University of Geneva.

### Data Processing

#### Alignment

FastQ files of the sequencing reads were aligned on the mouse reference genome NCBI Build GRCm38/mm10 using the Burrows-Wheeler Alignment tool v.0.7.17. The resulting SAM files were converted to BAM files keeping only the uniquely mapped reads. Samtools v1.9 was used to sort, remove the PCR duplicates and index the BAM file (https://arxiv.org/abs/1303.3997).

#### BigWig tracks

The processed BAM files have been transformed to coverage bigWig tracks using the bamCoverage function of deeptools v.3.5.1. The coverage was calculated as the normalized number of reads per bins per million mapped reads (BPM) with minimum mapping quality of 30 (http://doi.org/10.1093/nar/gkw257).

#### Replication timing domains

For the characterization of the replication timing domains of mESCs, MEFs and MSC the processed BAM files of Repli-seq experiment were given as input to the Repliscan tool (Zynda et al., 2017) using a window size of 10kb.

#### Peak calling

For the mapping of early IZs, we used an EdU-seq dataset from mESCs that were synchronized with nocodazole and MSO, released from the block in media containing EdU and hydroxyurea, and collected 2 h later (Fig. 1e, f: track i). Mid and late IZs were mapped using an EdU-seq dataset from mESCs 8 h after release from a MSO; the cells were treated with a CDC7 inhibitor between 3 and 6 h after release from the MSO and with hydroxyurea for 1.5 h prior to harvesting (Fig. 1e, f: track k). An in-house bash script was developed to call the origins in early, mid and late replicated domains (as identified from Repli-seq experiments). For the different samples, we divided the genome in early, mid and late domains according to the Repli-seq experiments. The chromosomes were split into 1kb bins and the EdU-seq signal was extracted from bigWig tracks using bigWigAverageOverBed (UCSC https://hgdownload.cse.ucsc.edu/admin/exe/linux.x86_64/). Given the variability in the distribution of signal among different RT domains (higher in early, lower in late RT domains), we established a signal cut-off for early, mid and late domains corresponding to 50%, 25% and 5% of the distribution of the highest signal bins, respectively, for each domain. Called regions were further refined in two consecutive rounds of genome binning (1kb) followed by signal extraction. Bins corresponding to the top 50% of the signal (median) were selected to define IZs. This method turned to be efficient for the identification of origins that tended to be clustered in short genomic areas, especially in the early-S replicating domains. Origins that overlapped with the genomic regions included in the ENCODE mm10 blacklist (regions with anomalous, unstructured, or high signal in next-generation sequencing experiments independent of cell line or experiment) were discarded as potential artifacts. The center of every origin was assessed by the refinepeak function of the macs3 algorithm using as input the coordinates of the called origins and the respective processed BAM file. The mid and late origins were shortlisted by discarding those that could potentially be confused with ongoing forks coming from earlier replicated domains. Given the presence of different cell populations (due to the nature of the bulk experiments) and to avoid false positive calls, we discarded those mid and late origins that mapped <500kb from neighbouring replication domains from which ongoing forks could come (for mid origins, early RT domains; for late origins, early and mid RT domains). The choice of 500kb was decided after calculating the fork speed from the EdU-seq experiments. See scheme in supplementary information. To create a random dataset of control regions for the signal analysis of mESC IZs, we first generated 10 million random positions of one base pair size using the mouse reference genome mm10 (chr1-X). Subsequently, these positions were intersected with the regions originated from the extension of the refined early, mid and late IZs for 50kb in both directions. The random positions that overlapped with the span of 50kb downstream - refined IZ center - 50kb upstream were discarded. Lastly, the non-overlapping random positions were shuffled and the first 1000 positions were selected as the final control dataset.

#### IZ S-phase plotting

The EdU BPM normalized signal corresponding to the center of every mESC Early-Mid-Late IZ was extracted for the time points 2 to 12 h from the mESC EdUseq timecourse experiments. The signal was transformed to log2+1 scale to avoid the log transformation error of zero values associated with the absence of replication signal of the IZs at specific timepoints. Every dot depicts the signal in the center of an IZ. Red lines indicate the median values of every group.

#### EdU-seq average signal

The EdU normalized signal (BPM) around every IZ (500 kb upstream and downstream) was extracted at 1 kb bin resolution. The values of each IZ group (ES -early S-, MS -mid S-, and LS -late S-) were averaged per bin position. For every group of IZs the bin with the lowest averaged BPM value was considered as background level and was hence subtracted from the averages of every bin within the same group.

#### Fork progression

To monitor fork progression, we used a subset of isolated IZs, which were called from the first fraction (G1) of the Repli-seq mESC experiments. We defined isolated IZs as those located at least 100 kb away from any other IZ in the same Repli-seq dataset. For these isolated IZs, we plotted the average EdU-seq signal from all the available time-course experiments (as seen in the figures). The centre value was then refined to 10 kb resolution through the refinepeak function of macs3 and used to extract the distance the fork travelled between the time points.

### Data and code availability

All the data produced in this study has been deposited in GEO database under accession numbers GSE271841, GSE271846, GSE271847. The bash script for peak calling can be downloaded from github (https://github.com/VSDionellis/my.scripts/blob/master/peak_calling.1kb.sh).

## ACKNOWLEDGEMENTS

We are grateful to David Suter, Cédric Deluz and Francesc Ventura for sharing cell lines and giving us technical support. We would like to acknowledge the help of Maëlys Alemany on cytometry analysis and Samia Barriot, Christine Seguin and Laurence Tropia for technical support. We thank the members of the Halazonetis laboratory for helpful discussions and suggestions. We would like also to thank the Flow Cytometry (Jean-Pierre Aubry-Lachainaye and Grégory Schneiter) and Genomics platforms (Mylène Docquier, Brice Petit, Christelle Barraclough and Didier Chollet) of the University of Geneva. This study was supported by grants from the European Commission (788681, REPLISTRESS), the Swiss National Science Foundation (212666) and the ACLON Foundation.

## AUTHORS CONTRIBUTION

Conceptualization and manuscript writing: E.R.-C. and T.D.H.

Experimental work: E.R.-C., L.B., S.G.N. and E.K.

Data processing: V.D.

Analysis of results: E.R.-C., V.D. and T.D.H. Funding acquisition: T.D.H.

## Supplementary Figures

**Supplementary Figure 1. Kinetics of progression through S phase.**

(**a**) EdU pulse chase experiment monitoring progression through S phase by flow cytometry. Asynchronous cells were pulsed with EdU for 30 minutes, washed and released into fresh media. Cells were collected at the indicated time points. Progression through S phase could be monitored for both the EdU-positive cells and the cells that were not in S phase during the EdU pulse (EdU-negative cells; oblique arrows at the 8 h time point). For mESCs, vertical arrows at the 9 and 12 h time points indicate cells that entered S phase of a second cell cycle.

(**b**, **c**) Genome-wide RT profile, as determined by Repli-seq. (**b**) Unsynchronized mESCs, MEFs and mMSCs were separated in fractions according to DNA content by flow sorting. (**c**) Repli-seq profiles of the indicated cell fractions. RT domains and annotated genes are indicated as in Fig. 1c. For mESCs, the RT domains determined from a published dataset (RT Zhao) ^17^ are also shown, revealing high similarity to the data obtained here.

(**c**) Genome-coverage by the RT domains in the different cell types and in (Zhao et al., 2020). ES, early S phase; ESMS, early to mid S transition; MS, mid S; MSLS, mid to late S transition; LS, late S; UND, undefined. Note that the analysis of IZs in this study did not include the ESMS, MSLS and UND regions, since these regions encompassed a small fraction of the genome and could not be unambiguously assigned to well-defined early, mid or late replicating domains.

(**d**) Clustering and correlation of genome-wide mESC EdU-seq and Repli-seq profiles. The data were obtained from the experiments outlined in Fig. 1b and Supplementary Fig. 1b.

**Supplementary Figure 2. IZ peak calling.**

(**a**) Workflow of IZ peak identification in mESC.

(**b**) Exemple of a region in chromosome 8 showing EdU-seq signal as in tracks i, e and k from Fig. 1e and f. The tracks below the EdU-seq show early, mid and late IZs before (preliminary) and after (definitive) refinement. The track above shows RT domains from the mESC Repli-seq experiments.

(**c**) Individual EdU signal plotting of every identified early, mid and late IZ at every time point of the EdU-seq time-course.

**Supplementary Figure 3. Lack of effect of increased concentration of nucleosides on RT.**

(**a**) Overlay of mESC Repli-seq profiles exploring the effect of adding nucleosides (0.5x Embryomax) to the tissue culture media. RT domains and annotated genes are indicated as in Fig. 1c.

(**b**) Overlay of mESC EdU-seq profiles exploring the effect of adding nucleosides (0.5x Embryomax) to the tissue culture media. RT domains and annotated genes are indicated as in Fig. 1c. Top: experimental outline.

(**c**) Clustering and correlation of genome-wide mESC Repli-seq profiles exploring the effect of adding nucleosides (Nuc, 0.5x Embryomax) to the tissue culture media. The data are derived from the experiment shown in panel a.

(**d**) Clustering and correlation of genome-wide mESC EdU-seq profiles exploring the effect of adding nucleosides (Nuc, 0.5x Embryomax) to the tissue culture media. The data are derived from the experiment shown in panel b.

**Supplementary Figure 4. Decreased nucleotide levels slow S phase progression without affecting relative RT.**

(**a**-**c**) Overlay of mESC EdU-seq profiles exploring the effect of adding HU to the tissue culture media. Three different genomic regions are shown. RT domains and annotated genes are indicated as in Fig. 1c.

(**a**) (**d**) Clustering and correlation of genome-wide mESC EdU-seq profiles exploring the effect of adding HU (100 μM) to the tissue culture media. The data are derived from the experiment shown in panels a-c.

(**e**-**g**) Average EdU-seq signal around early, mid and late IZs from control and HU-treated mESCs harvested 6 and 10 h after MSO, respectively. The heatmaps below indicate the EdU-seq signal at each IZ. The data are derived from the experiment shown in panels a-c.

**Supplementary Figure 5. Open chromatin marks and recruitment of pluripotency factors at IZs.** EdU-seq profiles of mESCs (from Fig. 1f) highlighting mid and late IZs (green and orange arrows, respectively) aligned to pluripotency factor and histone mark ChIP-seq profiles over a representative genomic region. RT domains and gene annotation are indicated as in Fig. 1c.

**Supplementary Figure 6. OCT4 enhances firing of select mid and late IZs.**

(**a**) Immunoblot showing effective downregulation of OCT4 levels in ZHBTc4 cells treated with doxycycline (Dox). Cells were harvested at the time of MSO or 2, 4 and 7 h later. The experimental outline is shown in Fig. 5b.

(**b**) Timing of entry into S phase, as monitored by flow cytometry analysis of the fraction of EdU-positive cells at the indicated time points after MSO. ZHBTc4 cells were treated as shown in the experimental outline of Fig. 5b.

(**c**) EdU-seq profiles of ZHBTc4 cells at the indicated times after MSO or after release from a thymidine block. Profiles with OCT4 expression turned ON or OFF (doxycycline added 12 h before MSO) are compared. Plots below the EdU-seq profiles: ATAC-seq profiles from asynchronous ZHBTc4 cells with OCT4 expression turned ON or OFF (Friman et al., 2019) and OCT4 ChIP-seq profile from asynchronous mESCs (Chronis et al., 2017). Select IZs mapping to mid and late replicating regions are indicated by green and orange arrows, respectively. RT domains and gene annotation are indicated as in Fig. 1c.

(**d**) Scatterplots of EdU-seq signals 4h after MSO at early, mid and late IZs of ZHBTc4 cells expressing (ON) versus not expressing (OFF) OCT4. OCT4 expression was suppressed by administering doxycycline 12 h before MSO. The EdU-seq signal for each IZ was computed as BPM in a region +/-5 kb around the centre of the IZ and transformed into log2. The diagonal blue lines indicate a hypothetical perfect correlation; the green lines show the trend of the real data; the angle between the lines was calculated to highlight the differences. All IZs with ON/OFF rates >2.5 are highlighted in red. Some late IZs are numbered for referencing.

(**e, f**) Scatterplots of ATAC-seq signals at early, mid and late IZs, plotted as in (Fig. 5e) (e) and RT domains (f). At most late IZs, there is a decrease in the ATAC-seq signal, when OCT4 expression was turned off. The ATAC-seq data are from (Friman et al., 2019).

## REFERENCES

Ahuja, A.K., Jodkowska, K., Teloni, F., Bizard, A.H., Zellweger, R., Herrador, R., Ortega, S., Hickson, I.D., Altmeyer, M., Mendez, J., Lopes, M., 2016. A short G1 phase imposes constitutive replication stress and fork remodelling in mouse embryonic stem cells. Nature Communications 7, 10660. 10.1038/ncomms10660

Apostolou, E., Hochedlinger, K., 2013. Chromatin dynamics during cellular reprogramming. Nature 502, 462–471. 10.1038/nature12749

Audit, B., Zaghloul, L., Vaillant, C., Chevereau, G., D’Aubenton-Carafa, Y., Thermes, C., Arneodo, A., 2009. Open chromatin encoded in DNA sequence is the signature of ‘master’ replication origins in human cells. Nucleic Acids Research 37, 6064–6075. 10.1093/NAR/GKP631

Becker, K.A., Stein, J.L., Lian, J.B., Van Wijnen, A.J., Stein, G.S., 2010. Human embryonic stem cells are pre-mitotically committed to self-renewal and acquire a lengthened G1 phase upon lineage programming. Journal of cellular physiology 222, 103. 10.1002/JCP.21925

Brossas, C., Valton, A.-L., Venev, S.V., Chilaka, S., Counillon, A., Laurent, M., Goncalves, C., Duriez, B., Picard, F., Dekker, J., Prioleau, M.-N., 2020. Clustering of strong replicators associated with active promoters is sufficient to establish an early-replicating domain. The EMBO Journal 39, e99520. 10.15252/EMBJ.201899520

Bulut-Karslioglu, A., Macrae, T.A., Oses-Prieto, J.A., Covarrubias, S., Percharde, M., Ku, G., Diaz, A., McManus, M.T., Burlingame, A.L., Ramalho-Santos, M., 2018. The Transcriptionally Permissive Chromatin State of Embryonic Stem Cells Is Acutely Tuned to Translational Output. Cell Stem Cell 22, 369–383.e8. 10.1016/j.stem.2018.02.004

Cadoret, J.-C., Meisch, F., Hassan-Zadeh, V., Luyten, I., Guillet, C., Duret, L., Quesneville, H., Prioleau, M.-N., 2008. Genome-wide studies highlight indirect links between human replication origins and gene regulation. Proceedings of the National Academy of Sciences 105, 15837–15842. 10.1073/pnas.0805208105

Cayrou, C., Coulombe, P., Vigneron, A., Stanojcic, S., Ganier, O., Peiffer, I., Rivals, E., Puy, A., Laurent-Chabalier, S., Desprat, R., Méchali, M., 2011. Genome-scale analysis of metazoan replication origins reveals their organization in specific but flexible sites defined by conserved features. Genome Research 21, 1438–1449. 10.1101/GR.121830.111

Chen, N., Buonomo, S.C.B., 2023. Three-dimensional nuclear organisation and the DNA replication timing program. Current Opinion in Structural Biology 83, 102704. 10.1016/J.SBI.2023.102704

Chen, T., Dent, S.Y.R., 2013. Chromatin modifiers and remodellers: regulators of cellular differentiation. Nature Reviews Genetics 2013 15:2 15, 93–106. 10.1038/nrg3607

Chronis, C., Fiziev, P., Papp, B., Butz, S., Bonora, G., Sabri, S., Ernst, J., Plath, K., 2017. Cooperative Binding of Transcription Factors Orchestrates Reprogramming. Cell 168, 442–459.e20. 10.1016/j.cell.2016.12.016

Coronado, D., Godet, M., Bourillot, P.Y., Tapponnier, Y., Bernat, A., Petit, M., Afanassieff, M., Markossian, S., Malashicheva, A., Iacone, R., Anastassiadis, K., Savatier, P., 2013. A short G1 phase is an intrinsic determinant of naïve embryonic stem cell pluripotency. Stem Cell Research 10, 118–131. 10.1016/j.scr.2012.10.004

Das, S.P., Borrman, T., Liu, V.W.T., Yang, S.C.H., Bechhoefer, J., Rhind, N., 2015. Replication timing is regulated by the number of MCMs loaded at origins. Genome Research 25, 1886–1892. 10.1101/gr.195305.115

Dellino, G.I., Cittaro, D., Piccioni, R., Luzi, L., Banfi, S., Segalla, S., Cesaroni, M., Mendoza-Maldonado, R., Giacca, M., Pelicci, P.G., 2013. Genome-wide mapping of human DNA-replication origins: Levels of transcription at ORC1 sites regulate origin selection and replication timing. Genome Research 23, 1–11. 10.1101/gr.142331.112

DePamphilis, M.L., 1993. How transcription factors regulate origins of DNA replication in eukaryotic cells. Trends in Cell Biology 3, 161–167. 10.1016/0962-8924(93)90137-P

Dequeker, B.J.H., Scherr, M.J., Brandão, H.B., Gassler, J., Powell, S., Gaspar, I., Flyamer, I.M., Lalic, A., Tang, W., Stocsits, R., Davidson, I.F., Peters, J.M., Duderstadt, K.E., Mirny, L.A., Tachibana, K., 2022. MCM complexes are barriers that restrict cohesin-mediated loop extrusion. Nature 606, 197–203. 10.1038/s41586-022-04730-0

Dileep, V., Gilbert, D.M., 2018. Single-cell replication profiling to measure stochastic variation in mammalian replication timing. Nature Communications 9, 1–8. 10.1038/s41467-017-02800-w

Dukaj, L., Rhind, N., 2021. The capacity of origins to load MCM establishes replication timing patterns. PLOS Genetics 17, e1009467. 10.1371/journal.pgen.1009467

Forey, R., Poveda, A., Sharma, S., Barthe, A., Padioleau, I., Renard, C., Lambert, R., Skrzypczak, M., Ginalski, K., Lengronne, A., Chabes, A., Pardo, B., Pasero, P., 2020. Mec1 Is Activated at the Onset of Normal S Phase by Low-dNTP Pools Impeding DNA Replication. Molecular Cell 78, 396–410.e4. 10.1016/j.molcel.2020.02.021

Foss, E.J., Sripathy, S., Gatbonton-Schwager, T., Kwak, H., Thiesen, A.H., Lao, U., Bedalov, A., 2021. Chromosomal Mcm2-7 distribution and the genome replication program in species from yeast to humans. PLOS Genetics 17, e1009714. 10.1371/JOURNAL.PGEN.1009714

Foti, R., Gnan, S., Cornacchia, D., Dileep, V., Bulut-Karslioglu, A., Diehl, S., Buness, A., Klein, F.A., Huber, W., Johnstone, E., Loos, R., Bertone, P., Gilbert, D.M., Manke, T., Jenuwein, T., Buonomo, S.C.B., 2016. Nuclear Architecture Organized by Rif1 Underpins the Replication-Timing Program. Molecular Cell 61, 260–273. 10.1016/j.molcel.2015.12.001

Friman, E.T., Deluz, C., Meireles-Filho, A.C.A., Govindan, S., Gardeux, V., Deplancke, B., Suter, D.M., 2019. Dynamic regulation of chromatin accessibility by pluripotency transcription factors across the cell cycle. eLife 8. 10.7554/eLife.50087

Gilbert, D.M., Takebayashi, S.I., Ryba, T., Lu, J., Pope, B.D., Wilson, K.A., Hiratani, I., 2010. Space and Time in the Nucleus: Developmental Control of Replication Timing and Chromosome Architecture. Cold Spring Harbor Symposia on Quantitative Biology 75, 143–153. 10.1101/sqb.2010.75.011

Gnan, S., Josephides, J.M., Wu, X., Spagnuolo, M., Saulebekova, D., Bohec, M., Dumont, M., Baudrin, L.G., Fachinetti, D., Baulande, S., Chen, C.-L., 2022. Kronos scRT: a uniform framework for single-cell replication timing analysis. Nature Communications 13, 2329. 10.1038/s41467-022-30043-x

Goldman, M.A., Holmquist, G.P., Gray, M.C., Caston, L.A., Nag, A., 1984. Replication Timing of Genes and Middle Repetitive Sequences. Science 224, 686–692. 10.1126/SCIENCE.6719109

Guilbaud, G., Murat, P., Wilkes, H.S., Lerner, L.K., Sale, J.E., Krude, T., 2022. Determination of human DNA replication origin position and efficiency reveals principles of initiation zone organisation. Nucleic Acids Research 50, 7436–7450. 10.1093/NAR/GKAC555

Heffernan, T.P., Ünsal-Kaçmaz, K., Heinloth, A.N., Simpson, D.A., Paules, R.S., Sancar, A., Cordeiro-Stone, M., Kaufmann, W.K., 2007. Cdc7-Dbf4 and the Human S Checkpoint Response to UVC. Journal of Biological Chemistry 282, 9458–9468. 10.1074/JBC.M611292200

Hiratani, I., Ryba, T., Itoh, M., Yokochi, T., Schwaiger, M., Chang, C.-W., Lyou, Y., Townes, T.M., Schübeler, D., Gilbert, D.M., 2008. Global Reorganization of Replication Domains During Embryonic Stem Cell Differentiation. PLoS Biology 6, e245. 10.1371/journal.pbio.0060245

Hu, Y., Stillman, B., 2023. Origins of DNA replication in eukaryotes. Molecular Cell 83, 352–372. 10.1016/j.molcel.2022.12.024

Hulke, M.L., Massey, D.J., Koren, A., 2020. Genomic methods for measuring DNA replication dynamics. Chromosome Research 28, 49–67. 10.1007/S10577-019-09624-Y/FIGURES/3

Jaksik, R., Wheeler, D.A., Kimmel, M., 2023. Detection and characterization of constitutive replication origins defined by DNA polymerase epsilon. BMC Biology 21, 1–18. 10.1186/S12915-023-01527-Z/TABLES/3

Juan, A.H., Wang, S., Ko, K.D., Zare, H., Tsai, P.F., Feng, X., Vivanco, K.O., Ascoli, A.M., Gutierrez-Cruz, G., Krebs, J., Sidoli, S., Knight, A.L., Pedersen, R.A., Garcia, B.A., Casellas, R., Zou, J., Sartorelli, V., 2016. Roles of H3K27me2 and H3K27me3 Examined during Fate Specification of Embryonic Stem Cells. Cell Reports 17, 1369–1382. 10.1016/j.celrep.2016.09.087

Katsuno, Y., Suzuki, A., Sugimura, K., Okumura, K., Zineldeen, D.H., Shimada, M., Niida, H., Mizuno, T., Hanaoka, F., Nakanishi, M., 2009. Cyclin A-Cdk1 regulates the origin firing program in mammalian cells. Proceedings of the National Academy of Sciences of the United States of America 106, 3184–9. 10.1073/pnas.0809350106

King, H.W., Klose, R.J., 2017. The pioneer factor OCT4 requires the chromatin remodeller BRG1 to support gene regulatory element function in mouse embryonic stem cells. eLife 6. 10.7554/ELIFE.22631

Klein, K.N., Zhao, P.A., Lyu, X., Sasaki, T., Bartlett, D.A., Singh, A.M., Tasan, I., Zhang, M., Watts, L.P., Hiraga, S., Natsume, T., Zhou, X., Baslan, T., Leung, D., Kanemaki, M.T., Donaldson, A.D., Zhao, H., Dalton, S., Corces, V.G., Gilbert, D.M., 2021. Replication timing maintains the global epigenetic state in human cells. Science 372, 371–378. 10.1126/science.aba5545

Knott, S.R.V., Peace, J.M., Ostrow, A.Z., Gan, Y., Rex, A.E., Viggiani, C.J., Tavaré, S., Aparicio, O.M., 2012. Forkhead Transcription Factors Establish Origin Timing and Long-Range Clustering in S. cerevisiae. Cell 148, 99–111. 10.1016/J.CELL.2011.12.012

Komamura-Kohno, Y., Karasawa-Shimizu, K., Saitoh, T., Sato, M., Hanaoka, F., Tanaka, S., Ishimi, Y., 2006. Site-specific phosphorylation of MCM4 during the cell cycle in mammalian cells. The FEBS Journal 273, 1224–1239. 10.1111/J.1742-4658.2006.05146.X

Le, T.M., Poddar, S., Capri, J.R., Abt, E.R., Kim, W., Wei, L., Uong, N.T., Cheng, C.M., Braas, D., Nikanjam, M., Rix, P., Merkurjev, D., Zaretsky, J., Kornblum, H.I., Ribas, A., Herschman, H.R., Whitelegge, J., Faull, K.F., Donahue, T.R., Czernin, J., Radu, C.G., 2017. ATR inhibition facilitates targeting of leukemia dependence on convergent nucleotide biosynthetic pathways. Nature Communications 2017 8:1 8, 1–14. 10.1038/s41467-017-00221-3

Li, R., 1999. Stimulation of DNA replication in Saccharomyces cerevisiae by a glutamine- and proline-rich transcriptional activation domain. Journal of Biological Chemistry 274, 30310–30314. 10.1074/jbc.274.42.30310

Li, V.C., Ballabeni, A., Kirschner, M.W., 2012. Gap 1 phase length and mouse embryonic stem cell self-renewal. Proceedings of the National Academy of Sciences of the United States of America 109, 12550–12555. 10.1073/pnas.1206740109

Lin, D.I., Aggarwal, P., Diehl, J.A., 2008. Phosphorylation of MCM3 on Ser-112 regulates its incorporation into the MCM2-7 complex. Proceedings of the National Academy of Sciences of the United States of America 105, 8079–8084. 10.1073/PNAS.0800077105/SUPPL_FILE/0800077105SI.PDF

Liu, L., Michowski, W., Kolodziejczyk, A., Sicinski, P., 2019. The cell cycle in stem cell proliferation, pluripotency and differentiation. Nature Cell Biology 2019 21:9 21, 1060–1067. 10.1038/s41556-019-0384-4

Liu, Q., Guntuku, S., Cui, X.S., Matsuoka, S., Cortez, D., Tamai, K., Luo, G., Carattini-Rivera, S., DeMayo, F., Bradley, A., Donehower, L.A., Elledge, S.J., 2000. Chk1 is an essential kinase that is regulated by Atr and required for the G2/M DNA damage checkpoint. Genes & Development 14, 1448–1459. 10.1101/GAD.14.12.1448

Liu, Y., Ai, C., Gan, T., Wu, J., Jiang, Y., Liu, X., Lu, R., Gao, N., Li, Q., Ji, X., Hu, J., 2021. Transcription shapes DNA replication initiation to preserve genome integrity. Genome Biology 22, 176. 10.1186/s13059-021-02390-3

Long, H., Zhang, L., Lv, M., Wen, Z., Zhang, W., Chen, X., Zhang, P., Li, T., Chang, L., Jin, C., Wu, G., Wang, X., Yang, F., Pei, J., Chen, P., Margueron, R., Deng, H., Zhu, M., Li, G., 2020. H2A.Z facilitates licensing and activation of early replication origins. Nature 577, 576–581. 10.1038/s41586-019-1877-9

Lyu, J., Shao, R., Kwong Yung, P.Y., Elsässer, S.J., 2022. Genome-wide mapping of G-quadruplex structures with CUT&Tag. Nucleic Acids Research 50, e13–e13. 10.1093/nar/gkab1073

Macheret, M., Halazonetis, T.D., 2019a. Monitoring early S-phase origin firing and replication fork movement by sequencing nascent DNA from synchronized cells. Nature Protocols 14, 51–67. 10.1038/s41596-018-0081-y

Macheret, M., Halazonetis, T.D., 2019b. Monitoring early S-phase origin firing and replication fork movement by sequencing nascent DNA from synchronized cells. Nature Protocols 14, 51–67. 10.1038/s41596-018-0081-y

Macheret, M., Halazonetis, T.D., 2018. Intragenic origins due to short G1 phases underlie oncogene-induced DNA replication stress. Nature 555, 112–116. 10.1038/nature25507

Marahrens, Y., Stillman, B., 1992. A Yeast Chromosomal Origin of DNA Replication Defined by Multiple Functional Elements. Science 255, 817–823. 10.1126/SCIENCE.1536007

Marchal, C., Sima, J., Gilbert, D.M., 2019a. Control of DNA replication timing in the 3D genome. Nature Reviews Molecular Cell Biology 20, 721–737. 10.1038/s41580-019-0162-y

Marchal, C., Sima, J., Gilbert, D.M., 2019b. Control of DNA replication timing in the 3D genome. Nature Reviews Molecular Cell Biology 20, 721–737. 10.1038/s41580-019-0162-y

Martin, M.M., Ryan, M., Kim, R.G., Zakas, A.L., Fu, H., Lin, C.M., Reinhold, W.C., Davis, S.R., Bilke, S., Liu, H., Doroshow, J.H., Reimers, M.A., Valenzuela, M.S., Pommier, Y., Meltzer, P.S., Aladjem, M.I., 2011. Genome-wide depletion of replication initiation events in highly transcribed regions. Genome Research 21, 1822–1832. 10.1101/gr.124644.111

Masai, H., Matsui, E., You, Z., Ishimi, Y., Tamai, K., Arai, K., 2000. Human Cdc7-related kinase complex. In vitro phosphorylation of MCM by concerted actions of Cdks and Cdc7 and that of a criticial threonine residue of Cdc7 by Cdks. Journal of Biological Chemistry 275, 29042–29052. 10.1074/jbc.m002713200

Mayran, A., Drouin, J., 2018. Pioneer transcription factors shape the epigenetic landscape. Journal of Biological Chemistry 293, 13795–13804. 10.1074/JBC.R117.001232

Michowski, W., Chick, J.M., Chu, C., Kolodziejczyk, A., Wang, Y., Suski, J.M., Abraham, B., Anders, L., Day, D., Dunkl, L.M., Li Cheong Man, M., Zhang, T., Laphanuwat, P., Bacon, N.A., Liu, L., Fassl, A., Sharma, S., Otto, T., Jecrois, E., Han, R., Sweeney, K.E., Marro, S., Wernig, M., Geng, Y., Moses, A., Li, C., Gygi, S.P., Young, R.A., Sicinski, P., 2020. Cdk1 Controls Global Epigenetic Landscape in Embryonic Stem Cells. Molecular Cell 78, 459–476.e13. 10.1016/J.MOLCEL.2020.03.010/ATTACHMENT/72C7CD46-C664-42BD-8A1C-9107679E7FAE/MMC9.XLSX

Moiseeva, T.N., Yin, Y., Calderon, M.J., Qian, C., Schamus-Haynes, S., Sugitani, N., Osmanbeyoglu, H.U., Rothenberg, E., Watkins, S.C., Bakkenist, C.J., 2019. An ATR and CHK1 kinase signaling mechanism that limits origin firing during unperturbed DNA replication. Proceedings of the National Academy of Sciences 116, 13374–13383. 10.1073/pnas.1903418116

Navarro, C., Lyu, J., Katsori, A.M., Caridha, R., Elsässer, S.J., 2020. An embryonic stem cell-specific heterochromatin state promotes core histone exchange in the absence of DNA accessibility. Nature Communications 11, 1–14. 10.1038/s41467-020-18863-1

Niwa, H., Miyazaki, J.I., Smith, A.G., 2000. Quantitative expression of Oct-3/4 defines differentiation, dedifferentiation or self-renewal of ES cells. Nature Genetics 2000 24:4 24, 372–376. 10.1038/74199

Ostrow, A.Z., Kalhora, R., Gana, Y., Villwocka, S.K., Linke, C., Barberisb, M., Chena, L., Aparicioa, O.M., 2017. Conserved forkhead dimerization motif controls DNA replication timing and spatial organization of chromosomes in S. cerevisiae. Proceedings of the National Academy of Sciences of the United States of America 114, E2411–E2419. 10.1073/PNAS.1612422114/SUPPL_FILE/PNAS.201612422SI.PDF

Pauklin, S., Vallier, L., 2013. The Cell-Cycle State of Stem Cells Determines Cell Fate Propensity. Cell 155, 135–147. 10.1016/J.CELL.2013.08.031

Peric-Hupkes, D., Meuleman, W., Pagie, L., Bruggeman, S.W.M., Solovei, I., Brugman, W., Gräf, S., Flicek, P., Kerkhoven, R.M., van Lohuizen, M., Reinders, M., Wessels, L., van Steensel, B., 2010. Molecular Maps of the Reorganization of Genome-Nuclear Lamina Interactions during Differentiation. Molecular Cell 38, 603–613. 10.1016/J.MOLCEL.2010.03.016

Petryk, N., Dalby, M., Wenger, A., Stromme, C.B., Strandsby, A., Andersson, R., Groth, A., 2018. MCM2 promotes symmetric inheritance of modified histones during DNA replication. Science 361, 1389–1392. 10.1126/science.aau0294

Petryk, N., Kahli, M., d’Aubenton-Carafa, Y., Jaszczyszyn, Y., Shen, Y., Silvain, M., Thermes, C., Chen, C.-L., Hyrien, O., 2016. Replication landscape of the human genome. Nat Commun 7, 10208. 10.1038/ncomms10208

Pope, B.D., Ryba, T., Dileep, V., Yue, F., Wu, W., Denas, O., Vera, D.L., Wang, Y., Hansen, R.S., Canfield, T.K., Thurman, R.E., Cheng, Y., Gülsoy, G., Dennis, J.H., Snyder, M.P., Stamatoyannopoulos, J.A., Taylor, J., Hardison, R.C., Kahveci, T., Ren, B., Gilbert, D.M., 2014. Topologically associating domains are stable units of replication-timing regulation. Nature 515, 402–405. 10.1038/nature13986

Prorok, P., Artufel, M., Aze, A., Coulombe, P., Peiffer, I., Lacroix, L., Guédin, A., Mergny, J.L., Damaschke, J., Schepers, A., Ballester, B., Méchali, M., 2019. Involvement of G-quadruplex regions in mammalian replication origin activity. Nature Communications 10, 1–16. 10.1038/s41467-019-11104-0

Raghuraman, M.K., Brewer, B.J., 2010. Molecular analysis of the replication program in unicellular model organisms. Chromosome Research 18, 19–34. 10.1007/S10577-009-9099-X/FIGURES/5

Rainey, M.D., Bennett, D., O’Dea, R., Zanchetta, M.E., Voisin, M., Seoighe, C., Santocanale, C., 2020. ATR Restrains DNA Synthesis and Mitotic Catastrophe in Response to CDC7 Inhibition. Cell Reports 32, 108096. 10.1016/j.celrep.2020.108096

Rausch, C., Weber, P., Prorok, P., Hörl, D., Maiser, A., Lehmkuhl, A., Chagin, V.O., Casas-Delucchi, C.S., Leonhardt, H., Cardoso, M.C., 2020. Developmental differences in genome replication program and origin activation. Nucleic Acids Research 48, 12751–12777. 10.1093/nar/gkaa1124

Rhind, N., Gilbert, D.M., 2013. DNA Replication Timing. Cold Spring Harb Perspect Biol 5, a010132. 10.1101/cshperspect.a010132

Rhind, N., Yang, S.C.-H., Bechhoefer, J., 2010. Reconciling stochastic origin firing with defined replication timing. Chromosome Research 18, 35–43. 10.1007/s10577-009-9093-3

Rivera-Mulia, J.C., Buckley, Q., Sasaki, T., Zimmerman, J., Didier, R.A., Nazor, K., Loring, J.F., Lian, Z., Weissman, S., Robins, A.J., Schulz, T.C., Menendez, L., Kulik, M.J., Dalton, S., Gabr, H., Kahveci, T., Gilbert, D.M., 2015. Dynamic changes in replication timing and gene expression during lineage specification of human pluripotent stem cells. Genome Research 25, 1091– 1103. 10.1101/gr.187989.114

Roccio, M., Schmitter, D., Knobloch, M., Okawa, Y., Sage, D., Lutolf, M.P., 2013. Predicting stem cell fate changes by differential cell cycle progression patterns. Development (Cambridge) 140, 459–470. 10.1242/dev.086215

Rossetti, G.G., Dommann, N., Karamichali, A., Dionellis, V.S., Asensio Aldave, A., Yarahmadov, T., Rodriguez-Carballo, E., Keogh, A., Candinas, D., Stroka, D., Halazonetis, T.D., 2024. In vivo DNA replication dynamics unveil aging-dependent replication stress. Cell 187, 6220–6234.e13. 10.1016/j.cell.2024.08.034

Saldivar, J.C., Cortez, D., Cimprich, K.A., 2017. The essential kinase ATR: ensuring faithful duplication of a challenging genome. Nature Reviews Molecular Cell Biology 2017 18:10 18, 622–636. 10.1038/nrm.2017.67

Schwaiger, M., Stadler, M.B., Bell, O., Kohler, H., Oakeley, E.J., Schübeler, D., 2009. Chromatin state marks cell-type- and gender-specific replication of the Drosophila genome. Genes & Development 23, 589–601. 10.1101/gad.511809

Sela, Y., Molotski, N., Golan, S., Itskovitz-Eldor, J., Soen, Y., 2012. Human Embryonic Stem Cells Exhibit Increased Propensity to Differentiate During the G1 Phase Prior to Phosphorylation of Retinoblastoma Protein. Stem Cells 30, 1097–1108. 10.1002/STEM.1078

Seller, C.A., O’Farrell, P.H., 2018. Rif1 prolongs the embryonic S phase at the Drosophila mid-blastula transition. PLoS Biology 16, e2005687. 10.1371/journal.pbio.2005687

Sequeira-Mendes, J., Díaz-Uriarte, R., Apedaile, A., Huntley, D., Brockdorff, N., Gómez, M., 2009. Transcription Initiation Activity Sets Replication Origin Efficiency in Mammalian Cells. PLoS Genetics 5, e1000446. 10.1371/journal.pgen.1000446

Sima, J., Chakraborty, A., Dileep, V., Michalski, M., Klein, K.N., Holcomb, N.P., Turner, J.L., Paulsen, M.T., Rivera-Mulia, J.C., Trevilla-Garcia, C., Bartlett, D.A., Zhao, P.A., Washburn, B.K., Nora, E.P., Kraft, K., Mundlos, S., Bruneau, B.G., Ljungman, M., Fraser, P., Ay, F., Gilbert, D.M., 2019. Identifying cis Elements for Spatiotemporal Control of Mammalian DNA Replication. Cell 176, 816–830.e18. 10.1016/j.cell.2018.11.036

Simoneau, A., Zou, L., 2021. An extending ATR–CHK1 circuitry: the replication stress response and beyond. Current Opinion in Genetics & Development 71, 92–98. 10.1016/J.GDE.2021.07.003

Smith, O.K., Kim, R., Fu, H., Martin, M.M., Lin, C.M., Utani, K., Zhang, Y., Marks, A.B., Lalande, M., Chamberlain, S., Libbrecht, M.W., Bouhassira, E.E., Ryan, M.C., Noble, W.S., Aladjem, M.I., 2016. Distinct epigenetic features of differentiation-regulated replication origins. Epigenetics and Chromatin 9, 18. 10.1186/s13072-016-0067-3

Sotiriou, S.K., Kamileri, I., Lugli, N., Evangelou, K., Da-Ré, C., Huber, F., Padayachy, L., Tardy, S., Nicati, N.L., Barriot, S., Ochs, F., Lukas, C., Lukas, J., Gorgoulis, V.G., Scapozza, L., Halazonetis, T.D., 2016. Mammalian RAD52 Functions in Break-Induced Replication Repair of Collapsed DNA Replication Forks. Mol Cell 64, 1127–1134. 10.1016/j.molcel.2016.10.038

Soufi, A., Donahue, G., Zaret, K.S., 2012. Facilitators and impediments of the pluripotency reprogramming factors’ initial engagement with the genome. Cell 151, 994–1004. 10.1016/j.cell.2012.09.045

Stewart-Morgan, K.R., Petryk, N., Groth, A., 2020. Chromatin replication and epigenetic cell memory. Nature Cell Biology 2020 22:4 22, 361–371. 10.1038/s41556-020-0487-y

Sugitani, N., Vendetti, F.P., Cipriano, A.J., Pandya, P., Deppas, J.J., Moiseeva, T.N., Schamus-Haynes, S., Wang, Y., Palmer, D., Osmanbeyoglu, H.U., Bostwick, A., Snyder, N.W., Gong, Y.N., Aird, K.M., Delgoffe, G.M., Beumer, J.H., Bakkenist, C.J., 2022. Thymidine rescues ATR kinase inhibitor-induced deoxyuridine contamination in genomic DNA, cell death, and interferon-α/β expression. Cell Reports 40. 10.1016/j.celrep.2022.111371

Suski, J.M., Ratnayeke, N., Braun, M., Zhang, T., Strmiska, V., Michowski, W., Can, G., Simoneau, A., Snioch, K., Cup, M., Sullivan, C.M., Wu, X., Nowacka, J., Branigan, T.B., Pack, L.R., DeCaprio, J.A., Geng, Y., Zou, L., Gygi, S.P., Walter, J.C., Meyer, T., Sicinski, P., 2022. CDC7-independent G1/S transition revealed by targeted protein degradation. Nature 605, 357–365. 10.1038/s41586-022-04698-x

Takahashi, S., Miura, H., Shibata, T., Nagao, K., Okumura, K., Ogata, M., Obuse, C., ichiro Takebayashi, S., Hiratani, I., 2019. Genome-wide stability of the DNA replication program in single mammalian cells. Nature Genetics 51, 529–540. 10.1038/s41588-019-0347-5

Tian, M., Wang, Z., Su, Z., Shibata, E., Shibata, Y., Dutta, A., Zang, C., 2023. Integrative analysis of DNA replication origins and ORC/MCM binding sites in human cells reveals a lack of overlap. eLife 12. 10.7554/ELIFE.89548

Turner, J.L., Hinojosa-Gonzalez, L., Sasaki, T., Vouzas, A., Soto, M.S., Chakraborty, A., Alexander, K.E., Fitch, C.A., Brown, A.N., Ay, F., Gilbert, D.M., 2023. Master transcription factor binding sites are necessary for early replication control element activity. 10.1101/2023.10.22.5634977

van den Berg, J., van Batenburg, V., Geisenberger, C., Tjeerdsma, R.B., de Jaime-Soguero, A., Acebrón, S.P., van Vugt, M.A.T.M., van Oudenaarden, A., 2024. Quantifying DNA replication speeds in single cells by scEdU-seq. Nature Methods 2024 21:7 21, 1175–1184. 10.1038/s41592-024-02308-4

Vassilev, L.T., Tovar, C., Chen, S., Knezevic, D., Zhao, X., Sun, H., Heimbrook, D.C., Chen, L., 2006. Selective small-molecule inhibitor reveals critical mitotic functions of human CDK1. Proceedings of the National Academy of Sciences of the United States of America 103, 10660–10665. 10.1073/PNAS.0600447103/SUPPL_FILE/00447MOVIE1.MOV

Wang, W., Klein, K.N., Proesmans, K., Yang, H., Marchal, C., Zhu, X., Borrman, T., Hastie, A., Weng, Z., Bechhoefer, J., Chen, C.-L., Gilbert, D.M., Rhind, N., 2021. Genome-wide mapping of human DNA replication by optical replication mapping supports a stochastic model of eukaryotic replication. Molecular Cell 81, 2975–2988.e6. 10.1016/j.molcel.2021.05.024

Wang, X.Q., Lo, C.M., Chen, L., Ngan, E.S.W., Xu, A., Poon, R.Y.C., 2016. CDK1-PDK1-PI3K/Akt signaling pathway regulates embryonic and induced pluripotency. Cell Death & Differentiation 2017 24:1 24, 38–48. 10.1038/cdd.2016.84

Wen, Z., Zhang, L., Ruan, H., Li, G., 2020. Histone variant H2A.Z regulates nucleosome unwrapping and CTCF binding in mouse ES cells. Nucleic acids research 48, 5939–5952. 10.1093/NAR/GKAA360

White, E.J., Emanuelsson, O., Scalzo, D., Royce, T., Kosak, S., Oakeley, E.J., Weissman, S., Gerstein, M., Groudine, M., Snyder, M., Schübeler, D., 2004. DNA replication-timing analysis of human chromosome 22 at high resolution and different developmental states. Proceedings of the National Academy of Sciences of the United States of America 101, 17771–17776. 10.1073/PNAS.0408170101/ASSET/903E917A-BC84-474C-859C-5112E57DADDD/ASSETS/GRAPHIC/ZPQ0510467600005.JPEG

Zegerman, P., Diffley, J.F.X., 2010. Checkpoint-dependent inhibition of DNA replication initiation by Sld3 and Dbf4 phosphorylation. Nature 2010 467:7314 467, 474–478. 10.1038/nature09373

Zhao, P.A., Sasaki, T., Gilbert, D.M., 2020. High-resolution Repli-Seq defines the temporal choreography of initiation, elongation and termination of replication in mammalian cells. Genome Biology 21, 76. 10.1186/s13059-020-01983-8

Zynda, G.J., Song, J., Concia, L., Wear, E.E., Hanley-Bowdoin, L., Thompson, W.F., Vaughn, M.W., 2017. Repliscan: a tool for classifying replication timing regions. BMC Bioinformatics 18, 362. 10.1186/s12859-017-1774-x

