## Supplementary Figures for "Pluripotency factors enhance the firing efficiency of late DNA replication origins in mouse embryonic stem cells"

#### Supplementary Figure 1

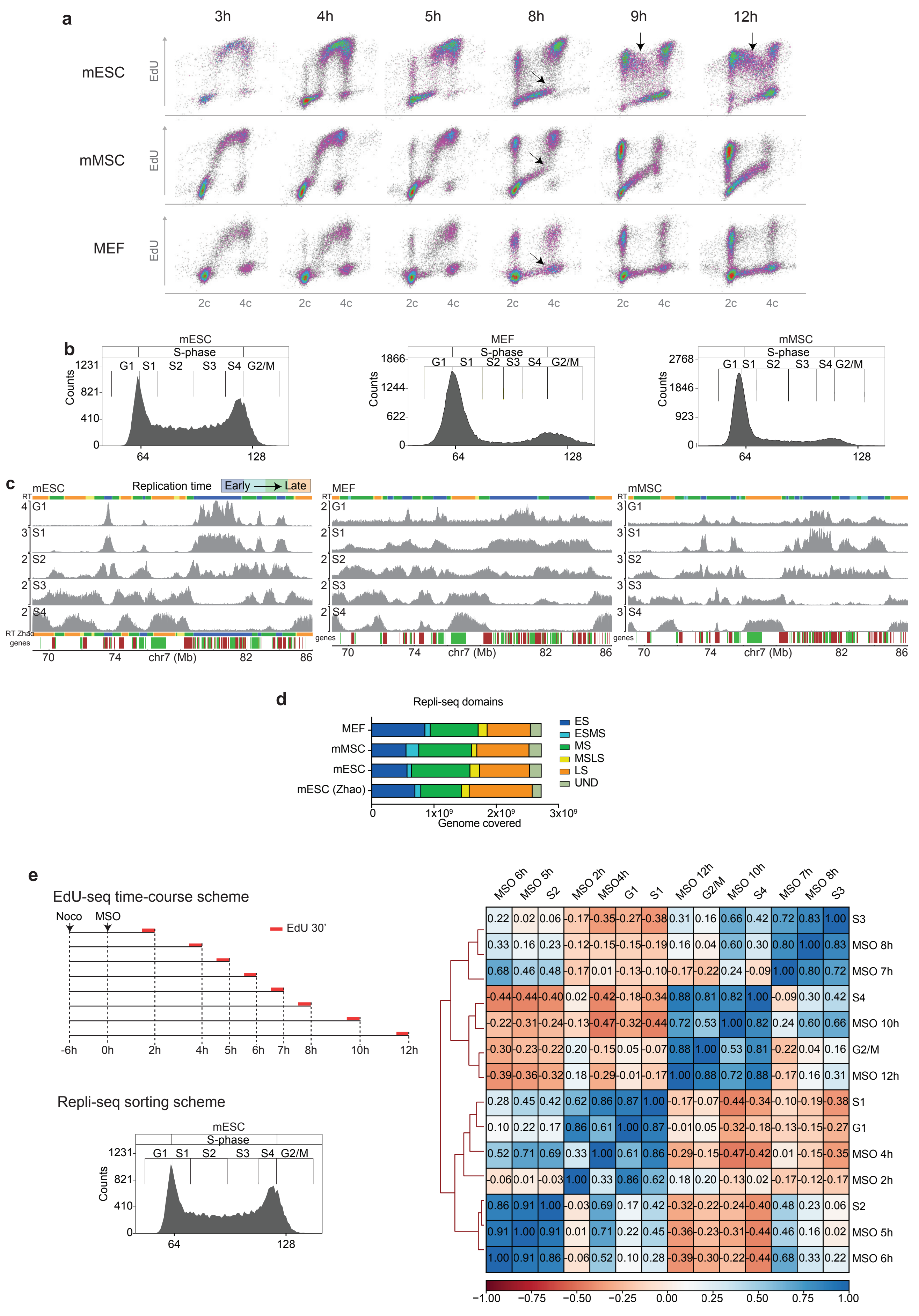

### Supplementary Figure 2

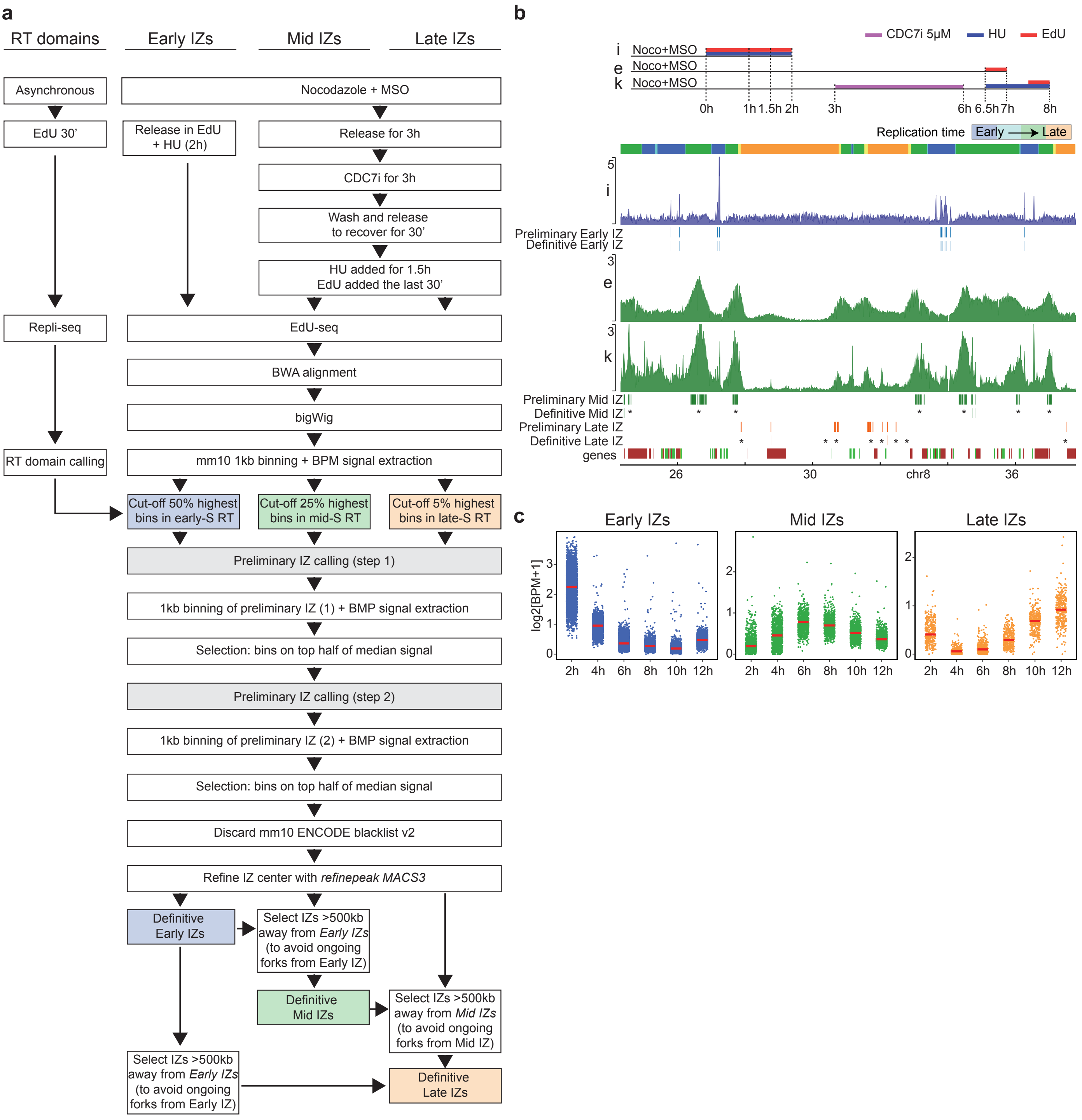

### Supplementary Figure 3

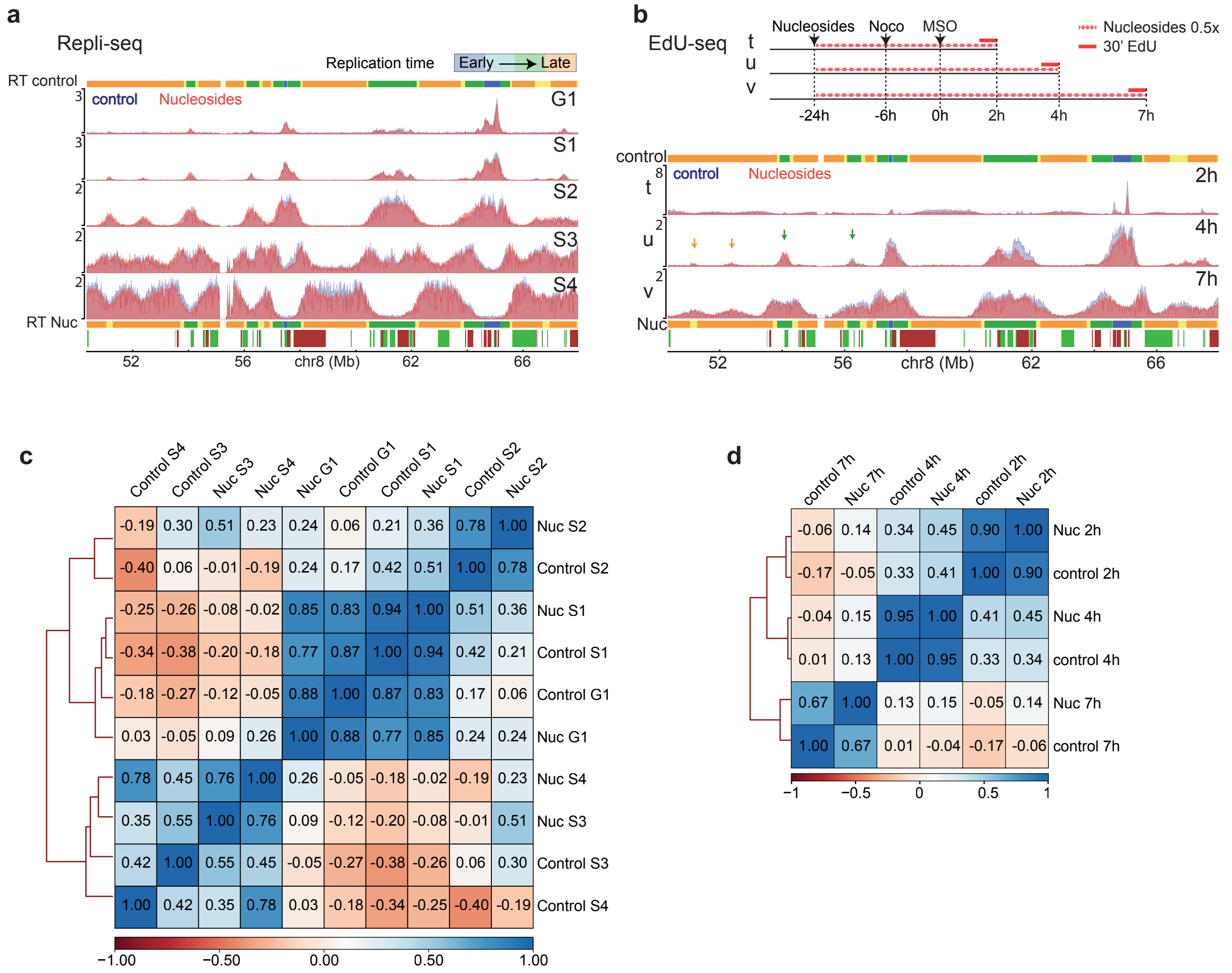

#### Supplementary Figure 4

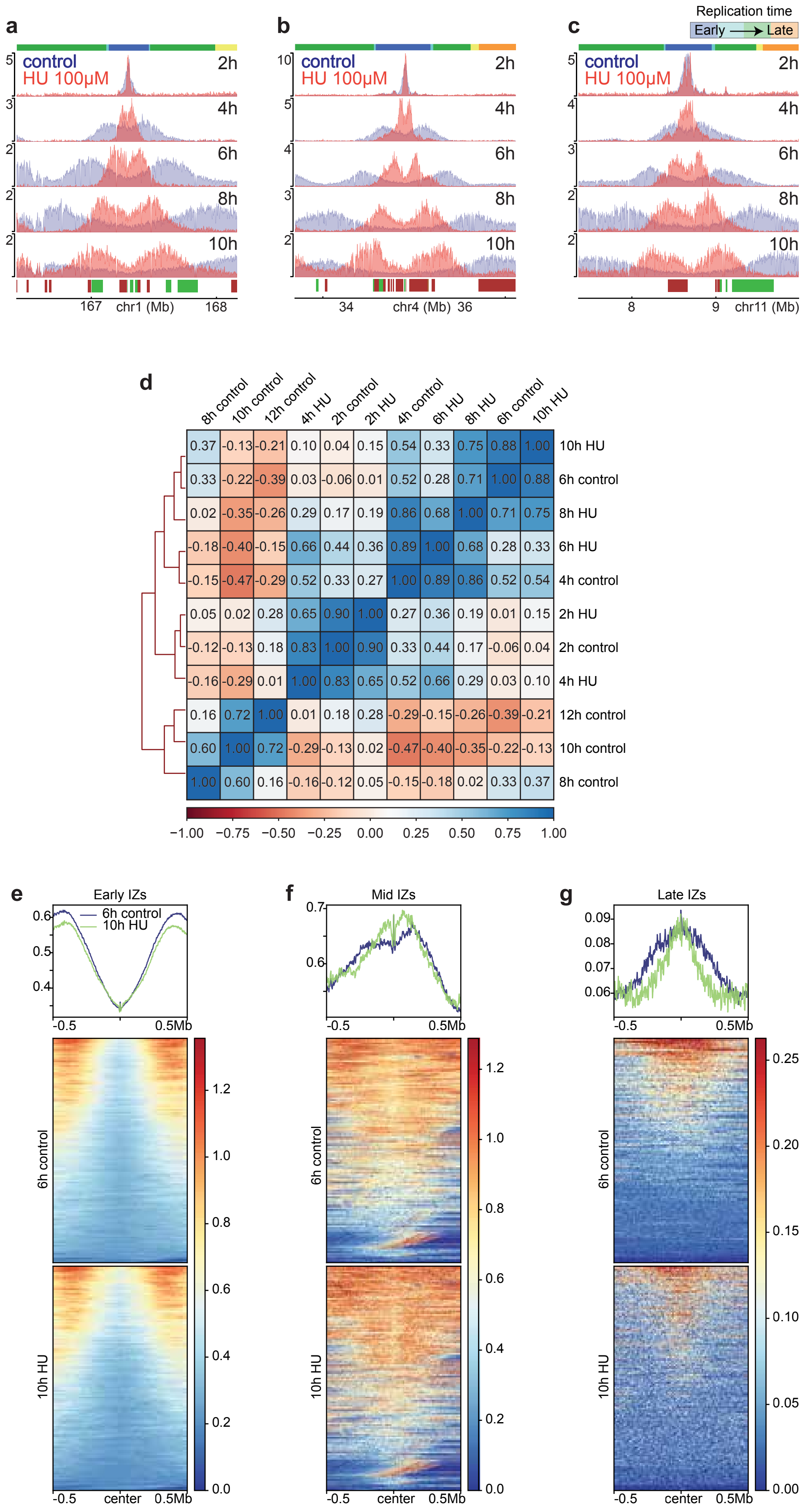

### Supplementary Figure 5

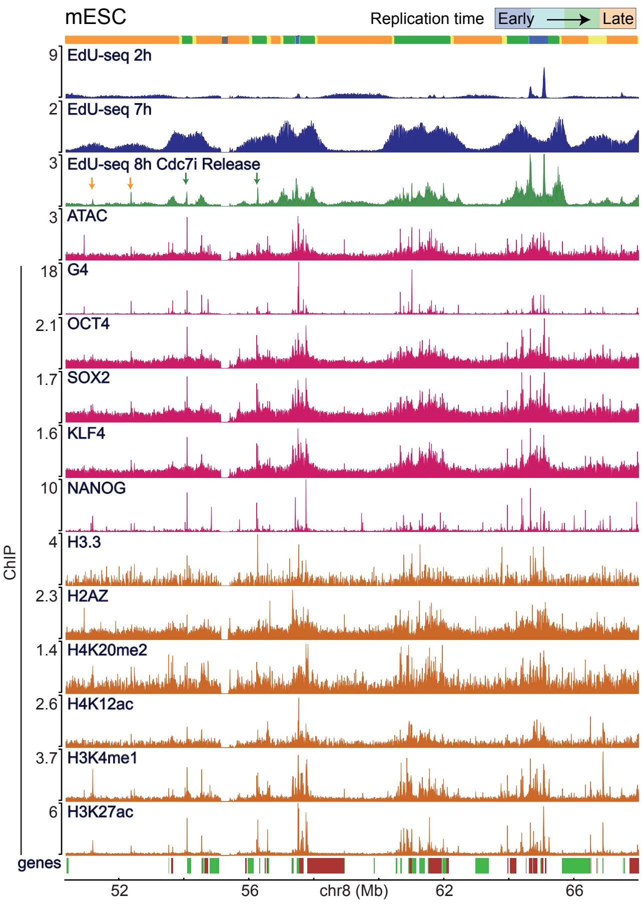

### Supplementary Figure 6

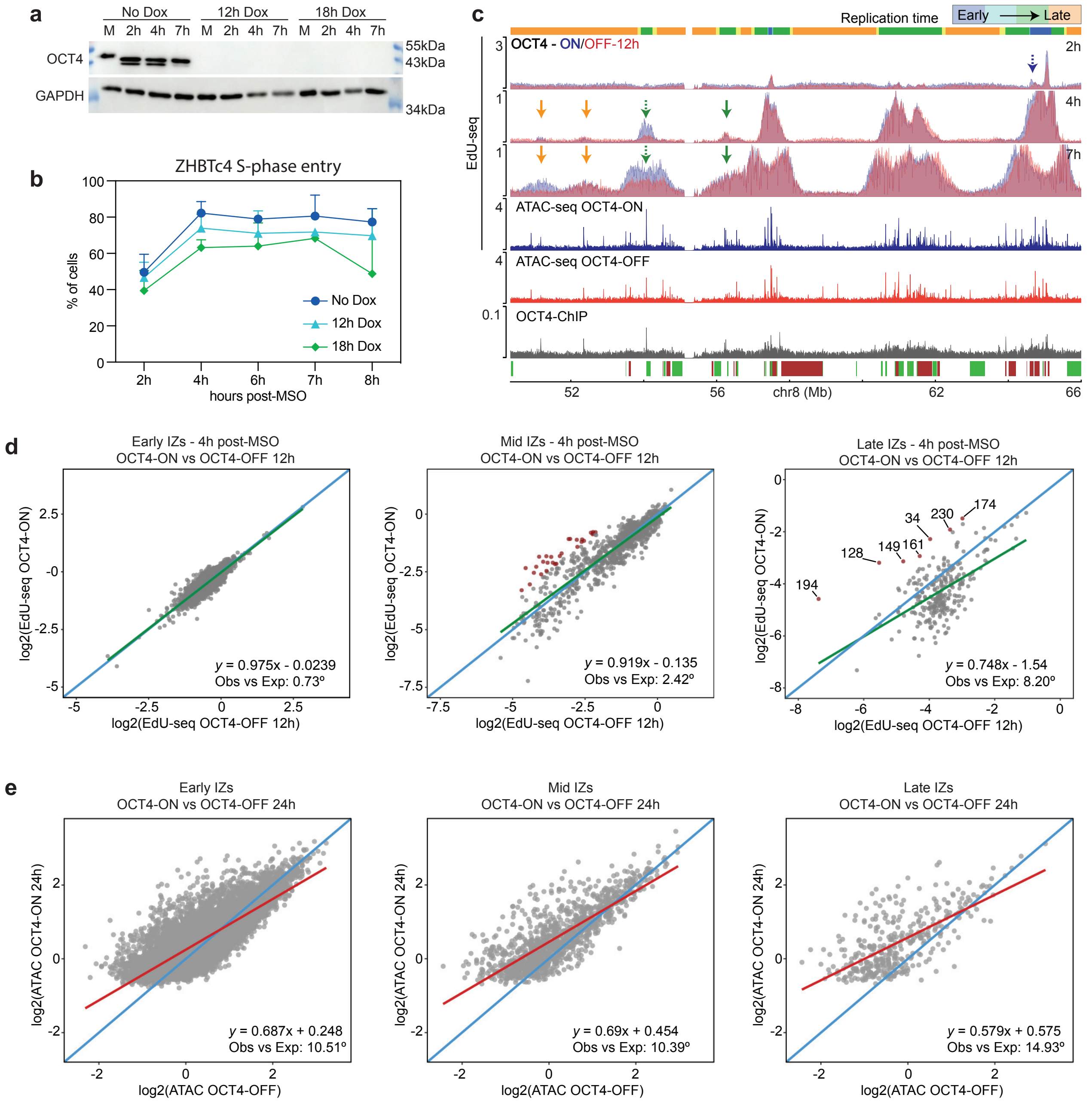
